# T-cell acute lymphoblastic leukemia progression is supported by inflammatory molecules including Hepatocyte Growth factor

**DOI:** 10.1101/2024.04.24.590927

**Authors:** Charly Le Maout, Lucine Fahy, Laurent Renou, Caroline Devanand, Charlotte Duwat, Vilma Barroca, Morgane Le Gall, Paola Ballerini, Arnaud Petit, Julien Calvo, Benjamin Uzan, Pflumio Françoise, Vanessa Petit

## Abstract

**Background:** T-cell acute lymphoblastic leukemia (T-ALL) is a malignant hematological disorder characterized by an increased proliferation of immature T lymphocytes precursors. T-ALL treatment includes chemotherapy with strong side effects, and patients that undergo relapse display poor prognosis. Although cell-intrinsic oncogenic pathways are well-studied, the tumor microenvironment, like inflammatory cellular and molecular components is less explored in T-ALL. We sought to determine the composition of the inflammatory microenvironment induced by T-ALL, and its role in T-ALL progression.

**Methods:** Two mouse T-ALL cell models were injected into immunocompetent mice. We used anti-Ly6G, and clodronate liposomes to suppress neutrophils and phagocytes, respectively. 5’- (N-ethylcarboxamido)adenosine (NECA), an agonist of adenosine receptors was used to decrease inflammatory molecules secretion.

**Findings:** We show that T-ALLs enhance blood neutrophils and resident monocytes, accompanied with a plasmatic acute secretion of inflammatory molecules. Depleting neutrophils or resident monocytes does not modulate plasmatic inflammatory molecule secretion and mice survival. However, inhibiting the secretion of inflammatory molecules by microenvironment with NECA diminishes T-ALL progression enhancing mouse survival. We uncovered Hepatocyte Growth Factor (HGF), T-ALL-driven and the most decreased molecule with NECA, as a potential therapeutic target in T-ALL.

**Interpretation:** Altogether, we identified a signature of inflammatory molecules that can potentially be involved in T-ALL evolution and uncovered HGF as a new potential therapeutic target.

**Fundings:** The work was supported by CEA, Inserm, Université Paris-Saclay and Université Paris-Cité, la Recherche contre le Cancer (ARC) and Hope of Princess Manon charity. The LSHL team is labellised by Ligue Nationale Contre le Cancer.

**Research in context:** *Evidence before this study:* T-cell acute lymphoblastic leukemia (T-ALL) is an aggressive and lethal hematologic malignancy accounting for about 15% of pediatric and 25% of adult ALL. T-ALL originates from a block of differentiation and uncontrolled proliferation of immature T cells. Current chemotherapies provide an overall 5 years survival higher than 90% in children and of about 50% in adults. Both pediatric and adult relapses have a very poor outcome with resistance to treatment. Therefore, the identification of molecular targets and the development of new specific therapies are major goals to improve treatment success, and one way to reach this goal is to have a better understanding the dialog between T-ALL cells and their microenvironment. Cellular and molecular actors in the microenvironment have been identified to impact several types of leukemia. Recently, the supportive role of myeloid cells has been described in T-ALL. Moreover, interactions between receptors and ligands such as DL1, IL-18, IL-7, IGF1 and CXCL12 sustain proliferation, survival or initiation of T-ALL. However, the composition and the contribution of the inflammatory microenvironment that may broadly help T-ALL progression still remains poorly explored.

*Added value of this study:* The study, utilizing NOTCH1 and TAL1/LMO1-driven mouse T-ALL models, reveals that T-ALL induces an inflammatory microenvironment characterized by increased levels of blood neutrophils, resident monocytes, and plasmatic inflammatory molecules. Targeting molecular microenvironment with the non-selective adenosine receptor agonist NECA drastically decreases T-ALL progression and prolongs mice survival. This study further identifies hepatocyte growth factor (HGF), a known regulator of proliferation and migration of tumor cells, as a putative supportive and targetable factor in T-ALL.

*Implications of all the available evidence:* In this study, evidence linking T-ALL and inflammatory microenvironment is provided. These data extend our understanding of the biological function of inflammatory microenvironment in T-ALL progression, and open to the targeting of the inflammatory microenvironment, and more specifically HGF/cMet signaling in T-ALL. Such targeted therapeutic approach could be added to current treatments to improve patient outcome.

## INTRODUCTION

T-cell acute lymphoblastic leukemia (T-ALL) is a hematologic malignancy that arises from the transformation of T-cell precursors in the thymus ^1^. Current treatments improve survival rates up to 90% for pediatric patients, whereas survival rates for adults remain around 50% ^2,3^. However, the intensified multiagent chemotherapy regimens are associated with significant long-term morbidity, including neurotoxicity, secondary cancers, and cardiopathies ^4,5^. Furthermore, frontline therapy refractory or relapsing T-ALL patients have poor outcomes and few alternative therapeutic options are available.

Earliest research on T-ALL has been focused on the identification of cell-intrinsic genetic alterations that promote tumor growth such as activating mutations in NOTCH1 pathway and aberrant expression of the transcription factors like T-cell acute lymphocytic leukemia 1 (TAL1/SCL) or LIM domain only 1/2 (LMO1/2) ^6–9^. Although oncogenic subgroups of T-ALL are identified, relapse and refractory patients can occur in various types of T-ALL subgroups supporting the idea of a predominant role of the niches to generate differential growth and treatment response of T-ALL cells in patients ^10,11^. Despite our deeper understanding of these cell-intrinsic alterations, we have yet to develop successful, possibly more targeted, therapeutic approaches to improve patient outcomes, suggesting the need to broaden our understanding of other factors affecting T-ALL.

Tumor progression is not only driven by cell-intrinsic alterations but also by dynamic interactions between cancer cells and the surrounding tumor microenvironment (TME). Notably, neither primary mouse nor human T-ALL cells survive when cultured in the absence of cytokines or cellular support suggesting that TME provides critical support for T-ALL growth ^12–14^. In addition to TME cells, molecular actors interact with leukemic blasts in a paracrine manner, including cytokines, chemokines or growth factors. Multiple cell types in the TME have been shown to support T-ALL survival and progression, including vascular endothelial cells through a C-X-C motif chemokine receptor 4 (CXCR4)-CXCL12 axis in the Bone Marrow (BM) ^15,16^, BM-derived stromal cells by pro-inflammatory cytokine IL-18 production ^17^ and thymic epithelial cells (TECs) through release of interleukin 7 (IL-7) ^18^ and expression of NOTCH1 ligands ^12^.

Biological features, such as inflammation can also participate in cancer progression, particularly in the case of hematological malignancies ^19^. The inflammatory microenvironment of cancers is a complex network in which cellular and molecular immune infiltration can contribute to the onset, the spread of cancerous cells, metastasis and the initiation of resistance mechanisms to anticancer therapies ^20,21^. Cytokines are essential for BM niche homeostasis, and the first studies on B-ALL reported a higher expression of these molecules when compared with healthy BM samples, suggesting an autocrine/paracrine regulation of leukemic cells and cytokines ^22^. More recently, studies have demonstrated that inflammation can drive progression of BCR-ABL1 B-ALL ^23^. The composition and the contribution of the inflammatory microenvironment in T-ALL still remains poorly explored. The deletion of stromal interaction molecule (STIM1/2) regulating store-operated Ca^2+^ entry (SOCE) mechanism improves T-ALL mouse survival in relation with reduced inflammation ^24^. Also, tumor-associated myeloid cells, in particular dendritic cells, support T-ALL growth via insulin-like growth factor 1 receptor (IGF1R) activation ^25^. These results suggest that inflammatory microenvironment including myeloid cells and soluble factors help T-ALL growth. Thus, T-ALL induced cellular and molecular inflammatory microenvironment response and its impact on leukemic progression still need better characterization.

In the present work, we investigated the role of the inflammatory microenvironment induced by mouse leukemia using NOTCH1 and TAL1/LMO1-driven T-ALL models. We found that T-ALL favors the recruitment of myeloid cells like neutrophils and resident monocytes accompanied with acute secretion of inflammatory molecules in the plasma. Depletion of specific myeloid cells was not sufficient to decrease inflammatory molecule secretion and increase T-ALL mice survival, even though leukemic burden could be impaired. Besides, inhibiting inflammatory molecule secretion in the plasma by a nonselective adenosine receptor agonist, 5’-(N-ethylcarboxamido)adenosine (NECA) treatment drastically decreased T-ALL progression and prolonged mouse survival. Our findings demonstrate the importance and characterize the composition of inflammatory microenvironment factor in T-ALL progression revealing HGF as a putative driver of T-ALL that can serve as opportune therapeutic target.

## METHODS

### Ethics statements

#### Mice

All C57BL/6 mice were housed and bred in CEA, Fontenay-aux-Roses, France, in a conventional animal facility (registration number C9203202) and handled in compliance with French Ministry of Research. Experimental procedures were performed in accordance with the European Community Council Directive (EC/2010/63) and were approved by the Ethics Committee (APAFIS#20682-2019051615369428 v2). All experiments were performed with 8 to 16 week-old-female mice.

#### Human T-ALL

7 primary T-ALL cell samples (hT-ALL) were collected by the Pediatric Hematology clinicians of Hôpital Armand Trousseau, Paris, France, after informed consent of the patients or the patient’s relatives in accordance with the Declaration of Helsinki and French ethics regulations. Prior to white blood cell purification, blood samples were centrifuged to harvest plasma samples that were stored at -80°C. Authorization KGM-13-16 was obtained from the ethics committee of Inserm (IORG0003254, FWA00005831). Patients’ characteristics are described in Supplementary Table S1.

### T-ALL mouse models

#### Notch1 induced T-ALL model (T-ALL#1)

T-ALL#1 cells were generated as described in ^9,26,27^. ICN1/T-ALL#1 containing intracellular domain of NOTCH1 (ICN1) construct followed by an IRES GFP reporter cells were kindly provided by Dr J Ghysdael, Institut Curie, Orsay. To perpetuate T-ALL#1, 1.5.10^4^ T-ALL#1 GFP^+^ cells were injected intravenously (i.v.) via retro-orbital injection into syngeneic immunocompetent wild type CD45.2 C57BL/6 littermates (weight, 18-25 g) after isoflurane anesthesia. The viability of T-ALL#1 cell suspension was assessed by flow cytometry with Zombie Aqua Viable dye (423102, BioLegend) prior to mouse injection. Mice were euthanized upon the detection of disease signs such as arched back, weight loss, disheveled hair, or at indicated time points by cervical dislocation. At sacrifice, up to 1ml of blood was recovered in Anticoagulant Citrate Dextrose (ACD-A, Macopharma); femurs were flushed in PBS; spleens and livers were weighed, crushed and cell suspensions were filtered with 80µm diameter nylon filters to remove debris. Red cells from all samples were lyzed with ACK Lysing Buffer (A10492-01, Gibco) following manufacturer’s guidelines and filtered with 80µm diameter nylon filters to remove debris. Plasma was recovered after centrifugation of blood samples at 5000 rpm for 5 minutes at 4°C,and supernatant aliquots with addition of protease inhibitors were frozen at -80°C.

#### TAL1/ LMO1 model (T-ALL#2)

C57BL/6J mice harboring either the *pSIL/SCL* mutation inducing overexpression of human TAL1 oncogene ^6^ or the *lck^Pr^ Ttg-1* resulting in the overexpression of human LMO1 oncogene ^28^ were bred in CEA mouse experimentation housing facility following ethic committee recommendation, in ventilated cages. Mice were kindly provided by the lab of Pr. V. Asnafi, Necker Hospital, Paris, France, after authorization by Dr PD Aplan, Center for Cancer Research, NCI, Bethesda, USA. Genotyping of the mice were done using PCR with primers in Supplementary Table S2. When mice developed T-ALL symptoms as described for T-ALL#1, they were sacrificed and their BM, spleen, thymus and lymph nodes were collected, followed by surface marker phenotyping to characterize the T cell differentiation origin of the T-ALL cells. Cells (named T-ALL#2) were frozen and conserved in 90% Fetal Calf Serum (FCS) and 10% DMSO solution (D5879, Sigma Aldrich), or injected into CD42.1.2 C57BL/6 recipient mice. At sacrifice, organs were collected and processed as described for TALL#1.

### In vivo treatments of mice

#### Neutrophils depletion

Neutrophil depletion was performed using 100µg intraperitoneal (i.p.) injection of either anti-Gr1 (clone RB6-8C5, Bio X Cell) or anti-Ly6G (clone 1A8, Bio X Cell) depleting antibodies or the equivalent quantity of isotype control rat, IgG2b (clone LTF-2, Bio X Cell, sham control) every 3 days, corresponding to a total of 3 injections/mouse, between day 7 and day 13 after T-ALL#1 injection. For the survival curve experiment, 100µg of anti-Ly6G depleting antibody was injected daily for 5 days between day 7 and day 11. Neutrophil depletion and T-ALL#1 progression were assessed at day 14 using Flow Cytometry.

#### NECA treatments

5’-(N-Ethylcarboxamido) adenosine (NECA) (E2387, Sigma Aldrich) adenosine receptor agonist was resuspended in DMSO at a concentration of 14mg/ml and aliquoted at -20°C following manufacturer’s recommendation. 1µg/20g of mice of NECA or the equivalent quantity of DMSO was injected via i.p. route (diluted in 100µl PBS) every 3 days for a total of 3 injections between day 7 and day 13 after the T-ALL#1 injection. For the survival curve protocol, a 4^th^ injection of NECA was done at day 15 in T-ALL#1 mice. Every mouse was sacrificed at day 14 for T-ALL#1. For T-ALL#2 model, NECA was injected 3 times a week from day 14 until day 35, providing a total of 10 injections. Mice were sacrificed at day 36.

#### Phagocytes depletion

Clodronate-liposomes and phosphate-buffered saline (PBS)-liposomes were purchased from LIPOSOMA (C-010). 200µl of a 5mg/ml liposome solution were injected i.p. every 3 days for a total of 3 injections between day 7 and day 13 after T-ALL#1 injection according to published study ^25^. Mice were either sacrificed at day 14 or followed up for survival.

Both treated and vehicle-treated mice in each experiment were not kept after day 14 for T-ALL#1 model and day 36 for T-ALL#2 model and blood, BM, spleen and liver were then harvested. Leukemic progression and modulation of immune cell populations were assessed on Canto II (Beckton Dickinson) or LSR II (Beckton Dickinson) flow cytometers.

### Secreted molecules level measurements

#### Mice cytokine array analysis

A pool of plasma from 2 to 4 mice was loaded on the membrane of the Proteome Profiler Mouse XL cytokine Array (ARY028, R&D systems) following manufacturer’s recommendations. Chemiluminescence signal was quantified using Gbox (Syngene) device. The analysis of the spots intensities and background noise removal was performed using ImageQuant TL software (Version 8.1, GE Healthcare). Each spot intensity was quantified as duplicates and the mean calculated. The relative levels were calculated dividing the intensity of the signal obtained for each tested molecule by the intensity of positive control pre-loaded spots. The fold change was calculated by dividing the relative quantification of each molecule in every tested pool of plasma from treated mice by the relative quantification of each molecule in the pooled plasma of control mice.

#### Human cytokine array analysis

The plasma of 4 T-ALL patients (age 6 to 14 years old) were collected by the Pediatric Hematology clinicians of Hôpital Armand Trousseau, Paris, France, provided informed consent by the patients or their relatives had been obtained in accordance with the declaration of Helsinki and French ethics regulation. Plasma from 2 healthy donors (age 25 and 27 years old) was obtained with informed consent. Each human plasma sample was loaded independently on a membrane of the Proteome Profiler Human XL Cytokine Array (ARY022B, R&D systems) following the manufacturer’s recommendations and background noise removal was performed identically.

#### Multiplex analysis

The Luminex Discovery assay Mouse premixed multi-analyte kit (LXSAMSM-03, R&D systems) for mouse plasma samples, and Luminex Discovery Assay Human premixed Multi-Analyte kit (LXSAHM, R&D systems) for human plasma were utilized delivering concentrations of murine or human HGF in ng/µl after 2-fold dilution of plasma samples following manufacturer’s recommendations. HGF quantification of all samples was independently acquired using Bio-Plex 200 system (LX10016187401, BioRad) device. The monthly validation was successfully achieved using the Bio-Plex kit 4.0 (171203001, BioRad) and the calibration was set using Bio-Plex Calibration kit (17120360, BioRad) prior to sample quantification. Data were acquired and analyzed using the Bio-Plex Manager software (Version 6.1.1, BioRad).

### In vitro co-culture of T-ALL cells

MS5 BM-derived stromal cells were cultured in αMEM (32561-029, Gibco) complemented with 10% heat-inactivated Fetal Bovine Serum (FBS) (F9665, Sigma Aldrich), 1% Penicillin Streptomycin (15140-122, Gibco) and 1% L-Glutamine (25030-024, Gibco). 1.10^5^ T-ALL#1 or T-ALL#2 cells were seeded on a confluent layer of MS5 cells in RPMI 1640 (31870-025, Gibco, Life Technology, France) complemented with 10% FBS (F9665, Sigma Aldrich), 1% Penicillin Streptomycin (15140-122, Gibco) and 1% L-Glutamine (25030-024, Gibco) in a 24-well plate (3524, Corning). Culture medium was complemented with NECA (0.1, 1 and 10 µM). Cell growth was assessed by counting T-ALL cells 2 and 5 days after culture initiation by flow cytometry. Culture medium was complemented with SU11274 (cMet phosphorylation inhibitor, SE-S1080, Selleck Chemicals) at 5 µM. Cell growth, Annexin V apoptosis and cell cycle analysis were assessed by flow cytometry 24 hours after culture initiation.

### Flow cytometry and cell sorting

Blood, BM and spleen cells were stained with indicated antibodies at 4°C in the dark for 30 minutes. Antibodies were diluted at 1:500. Flow cytometry antibodies were listed in Supplementary Table S3. Following staining, cells were washed in PBS and resuspended in 300µl PBS. Cells were characterized using Canto II or LSR II flow cytometers (both BD Bioscience).

*Cell sorting* was performed with BM cells stained with anti-CD45 APC antibody (clone 30-F11, eBioscience) and dead cells were removed using HOECHST staining. HOECHST^-^CD45^+^ GFP^+^ cells were sorted using Aria or Influx cell sorting cytometers (both BD Biosystem).

*Cell cycle analysis* was done using 1-2.10^6^ of T-ALL#1 cells from BM, spleen and *in vitro* cultured cells, stained with anti-CD4 PerCP/Cyanine 5.5 (clone GK1.5, eBioscience) and anti-CD8a APC (clone 53-6.7, BD pharmingen) for 40 minutes at 4°C. Following surface marker staining, cells were washed and permeabilized using BD Cytofix/CytopermTM (BD Biosciences) according to the manufacturer’s protocol. Cells were then incubated with anti-Ki67 PE antibody (51-36525X, BD Bioscience) or with isotype PE (51-35405X, BD Bioscience) and with 40µg/ml HOECHST 33342 (H3570, Invitrogen) and cell cycle entry/ DNA content were measured.

*Apoptosis levels* were quantified in T-ALL#1 BM, spleen and *in vitro* cultured cells. Cell suspension was first washed with Annexin V binding buffer (51-66121E, BD Pharmingen) according to manufacturer’s protocol, then incubated with APC-Annexin V (640920, BioLegend) for 15 minutes at room temperature (RT) and extemporaneously with HOECHST 33258; 1,25 µg/ml final concentration (H3569, Invitrogen).

*cMet phosphorylation levels* were assessed by flow cytometry. First surface staining was performed using anti-CD4 PerCP/Cyanine 5.5 (clone GK1.5, eBioscience) and anti-CD8a APC (clone 53-6.7, BD pharmingen) for 30 minutes at 4°C. Cells were washed with cold PBS and resuspended in 70% ethanol while being smoothly vortexed, and maintained on ice for 30 minutes to insure proper permeabilization. Cell suspensions were then washed twice with cold PBS BSA 1% and stained overnight with 5µl of anti-phospho-cMet APC (clone MetY12341235-6F11, Thermofisher) following manufacturer’s recommendations or the equivalent of isotype control antibody (clone RIC-001-APC, Alomone labs). Cells were then washed twice with PBS BSA1% at 4°C and resuspended in 200µl cold PBS extemporaneously.

### Microarray analysis

Up to 2.10^6^ T-ALL#1 GFP^+^ cells were sorted from femurs of NECA or vehicle-treated T-ALL mice using the Influx cell sorter and mRNA was extracted directly after sorting using the RNeasy Plus Mini kit (74134, QIAGEN). The quality of the RNA was acquired using the RNA 6000 Pico Labchip Kit (5067-1513, Agilent) providing RNA Integrity Number (RIN) of 10 for each sample used for the microarray. Each sample RNA was then processed to micro-array analysis using GeneChip Whole Transcript Pico Reagent Kit expression array (902622, ThermoFisher) for amplification and Biotin incorporation and was further loaded onto Clariom D array Mouse Cartridge (520851, Applied Biosystem) for gene expression assessment. Data were then analyzed by Transcriptome Analysis Console (TAC) 4.0. All transcripts were submitted to Gene Set Enrichment Analysis (GSEA) using GSEA v4.0.3 software (https://www.gsea-msigdb.org) on Hallmark gene sets (MSigDB v7.5.1). Were considered as DEGs (differential expressed genes), transcripts with both False Discovery Rates (FDR) p-value <0.05 and |fold change|>2. The 268 selected DEGs, 170 upregulated and 98 downregulated in NECA compared to vehicle-injected mice, were analyzed using Ingenuity Pathway Analysis (IPA, QIAGEN Inc, version 70750971). Significantly over-represented canonical pathways were identified using a right-tailed Fisher’s Exact test that calculates an overlap Benjamini-Hochberg adjusted p-value (B-H p-value). The activation status of each pathway was also predicted using z-score calculation. This metric assessed the activation (positive z-score) or repression (negative z-score) of each pathway.

### Capillary electrophoresis immunoassay (Simple Western)

T-ALL#1 cells were maintained in coculture with MS5 that were eliminated prior to protein extraction. Total proteins from T-ALL#1 cells were extracted with a RIPA buffer containing a cocktail of 1Xprotease inhibitors (11836145001, Roche) and 1X phosphatase inhibitors (P2850; P5726, Sigma-Aldrich). After 5 min denaturation at 95°C in the presence of 5X Fluorescent Master Mix (PS-FL01-8, ProteinSimple), samples were assayed on a ProteinSimple Wes automated capillary-based electrophoresis instrument with Wes Separation Module protocol (ProteinSimple). The 12-230 kDa Separation Module 8 x 25 capillary cartridges (SM-W004, ProteinSimple) and the Anti-Rabbit Detection Module (DM-001, ProteinSimple) were used. Proteins were detected using rabbit primary antibodies listed in Supplementary Table S4. Results (peak area) were analyzed using Compass for SW software v5.0.1.

### Statistical analysis

Statistical comparisons were performed with Mann-Whitney non-parametric test, in accordance with data distribution. Mouse survival curves were obtained using the Kaplan-Meier statistics and log-rank Mantel-Cox statistical test was applied comparing treated or untreated conditions. Correlation analysis was performed using Spearman’s r correlation coefficient in accordance with data distribution. All statistical analyses were performed with GraphPad Prism 8.0 (GraphPad Software, CA).

### Role of funders

The founders played no roles in the study design, data collection, data analysis, interpretation, or the writing of the manuscript.

## RESULTS

### T-ALL increases blood neutrophils and resident monocytes correlated with high inflammatory molecules in the plasma

Previous studies have shown that even though leukemic cells become the most dominant cell population in the hematopoietic organs as leukemia develops, a supportive microenvironment is important for its establishment and rapid dissemination ^29^. Our goal being to better characterize the impact of the microenvironment in T-ALL progression, we first used a NOTCH1-induced mouse T-ALL model (further noted as T-ALL#1), NOTCH1 being a major actor in T-ALL pathology ^9^. In this model, intracellular NOTCH1 (ICN1), the active form of NOTCH1is constitutively expressed in T cell progenitors through a MIGR retrovirus vector that includes a GFP tracer ^26^. Phenotypic analysis showed that GFP^+^ T-ALL#1 express mature cortical T lymphoid progenitor markers CD4^+^CD8^+^TCR-β^+^CD3^+^ as described previously ^30^ (Fig. 1a). T-ALL#1 cells are detectable in the blood, BM and spleen 7 days after the injection of T-ALL#1 cells in non-irradiated C57BL/6 wild type (WT) mice and remain low until day 12. The highest levels of leukemia are observed at day 12 triggering a 14 days median survival time of leukemic mice (Fig. 1b and Supplementary Fig. S1a). Survival curves and dynamic distribution of T-ALL cells suggested that day 7 and 14 were interesting time points to define early and late stages of leukemia, respectively. At 14 days post T-ALL#1 mouse transplant, a 2.3- and 1.6-fold increase of spleen and liver weight was observed respectively, in comparison with control mice (Fig. 1c-d). Representative size of spleen and liver are illustrated in Fig. 1c-d (right panels). Extramedullary organ infiltration is a common clinical feature of leukemia and is observed in many T-ALL models ^31^. Because immune cells are part of the leukemic cell microenvironment in those organs, we followed the evolution of several immune cell populations among the normal cells (*i.e.* GFP^-^ cells) including neutrophils (CD11b^+^ Ly6G^+^), inflammatory monocytes (CD11b^+^ Ly6C^+^ CD115^+^), resident monocytes (CD11b^+^ Ly6C^+^ CD115^+^) and lymphoid cells including B lymphocytes (CD11b^-^ B220^+^) and T lymphocytes (CD11b^-^ CD3^+^) (Supplementary Fig. S1b). We observed that the progression of T-ALL was only correlated with the increase of blood neutrophils and resident monocytes with a 3.4-fold more neutrophils and 2.0-fold more resident monocytes at day 14 in case of T-ALL development (Fig. 1e-g and Supplementary Fig. S1c-d).

**Fig. 1:**
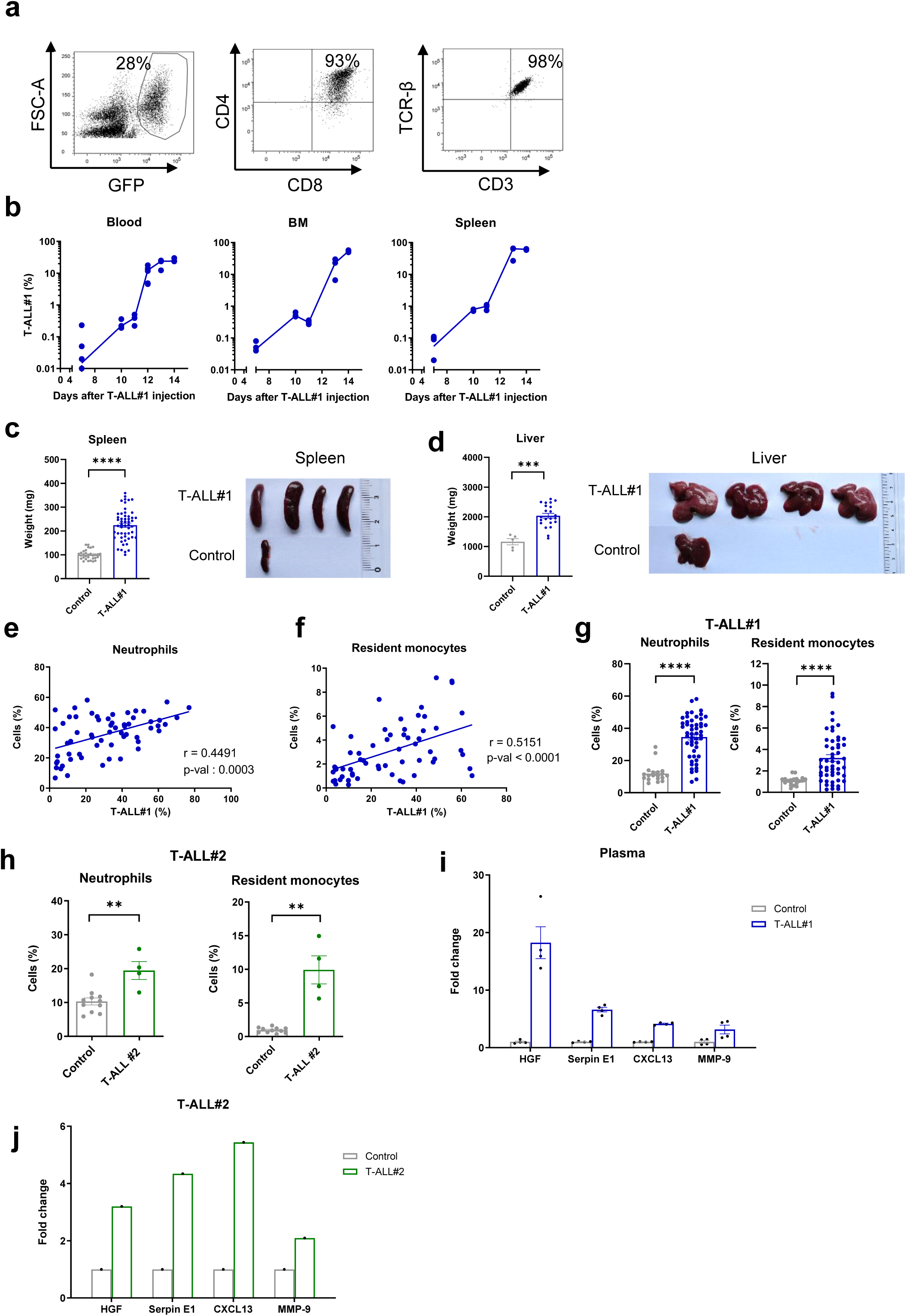
Mouse syngenic models of T-ALL increase blood neutrophils, resident monocytes and inflammatory molecule secretion. (a) Flow cytometry analysis of T-ALL#1 cells recovered from mouse BM at sacrifice. b) Kinetic evolution of the T-ALL#1 percent in the blood, BM and spleen at day 7, 10, 11, 13 and 14 after T-ALL#1 injection (log scale). n=3 mice/time point. (c-d) Weights of the spleen (c) and liver (d) and their representative size and morphology 14 days after T-ALL#1 injection compared to control mice. (31 spleens and 5 livers of control mice, 53 spleens and 22 livers of T-ALL#1 mice). (e-f) Correlation between the percent of blood neutrophils (e) or resident monocytes (f) and the percent of blood T-ALL#1 cells. Shown are results from 60 mice. Spearman’s r correlation test was applied. (g) Percent of blood neutrophils (left) and resident monocytes (right) in T-ALL#1 injected mice compared to control mice at day 14 of 19 control and 51 T-ALL#1 mice. (h) Percent of blood neutrophils (left) and resident monocytes (right) in T-ALL#2 (4 mice) injected mice compared to control mice (8 mice) (day 34). (i-j) Fold change of mouse HGF, Serpin E1, CXCL13 and MMP-9 levels identified by cytokine array in plasma samples after T-ALL#1 injection (i: day 14) or T-ALL#2 injection (j: day 36) compared to control plasma. Each dot represents one cytokine array experiment. 2 plasma samples were pooled for controls in 2 independent experiments. 3 or 4 plasma samples were pooled for T-ALL#1 in 4 independent experiments and 2 plasma were pooled for T-ALL#2 in 1 experiment. Data in c-d and g-i represent means ± SEM. **p<0.01, ***p<0.001, ****p<0.0001 (two-tailed Mann-Whitney U test).

To verify that these observations were independent of *ICN1*-T-ALL model, we performed identical experiments with another T-ALL model in which TAL1 and LMO1 oncogenes were combined ^6^. Mice developed T-ALL with CD4^-^CD8^-^CD44^-^CD25^+^ immature progenitor phenotype (further entitled T-ALL #2) (Supplementary Fig. S1e). The spreading of T-ALL#2 cells in several hematopoietic organs was observed 34 days post injection when T-ALL#2 cells were injected in CD45.1.2+ non irradiated C57BL/6 recipient mice (Supplementary Fig. S1f). Similarly to T-ALL#1, T-ALL#2 infiltrated mice exhibited a 2-fold and 10-fold increase of blood neutrophils and resident monocytes respectively (Fig. 1h). The observations made with these two T-ALL models suggest that T-ALL induces a cellular inflammatory microenvironment with increased neutrophils and resident monocytes in the peripheral blood.

To further characterize the inflammatory response associated with T-ALL, we tested the presence of inflammatory molecules in the plasma with relative quantification cytokine array. As shown in Fig. 1i-j, an increase of different class of inflammatory molecules was observed in the plasma of both T-ALL models such as growth factors, cytokines, chemokines and metalloproteases (Supplementary Fig. S1g-h). Notably, enhanced and significant levels of plasmatic hepatocyte growth factor (HGF) were observed in T-ALL#1 mice (20-fold, most increased molecule) and in T-ALL#2 mice (3-fold) as well as a significant increase of Serpin E1, CXCL13 and Matrix metallopeptidase 9 (MMP-9) compared to healthy mice (Fig. 1i-j).

Altogether, these results describe actors of the blood microenvironment that are stimulated and may interact with T-ALL cells, thus providing a favorable context to leukemic expansion. This environment includes increased blood neutrophils, resident monocytes and inflammatory molecules.

### T-ALL progression is independent of neutrophils *in vivo*

Since the increase of blood neutrophils is correlated with T-ALL growth *in vivo*, we investigated whether neutrophils could contribute to generalized inflammation and in turn, support T-ALL progression. To test this hypothesis, we assessed leukemic invasion in T-ALL#1 mice treated with anti-Gr1 antibody to deplete neutrophils. Anti-Gr1 recognizes Ly6G, the most widely used marker of mouse neutrophils, and Ly6C expressed on inflammatory monocytes ^32^. When blood T-ALL burden reached 0.1-1%, anti-Gr1 was administrated to half of the cohort with 3 intraperitoneal injections from day 7 to day 13 and T-ALL burden was followed up (Fig. 2a). As expected, data extracted from flow cytometry analysis indicated a selective decrease of myeloid cells mostly in blood and spleen in anti-Gr1-treated mice (Fig. 2b and Supplementary Fig. S2a-b). Despite this reduction, no difference in T-ALL burden in the BM and even an increase was noted in blood and spleen in percent and in absolute number (Fig. 2c and Supplementary Fig. S2a-b).

**Fig. 2:**
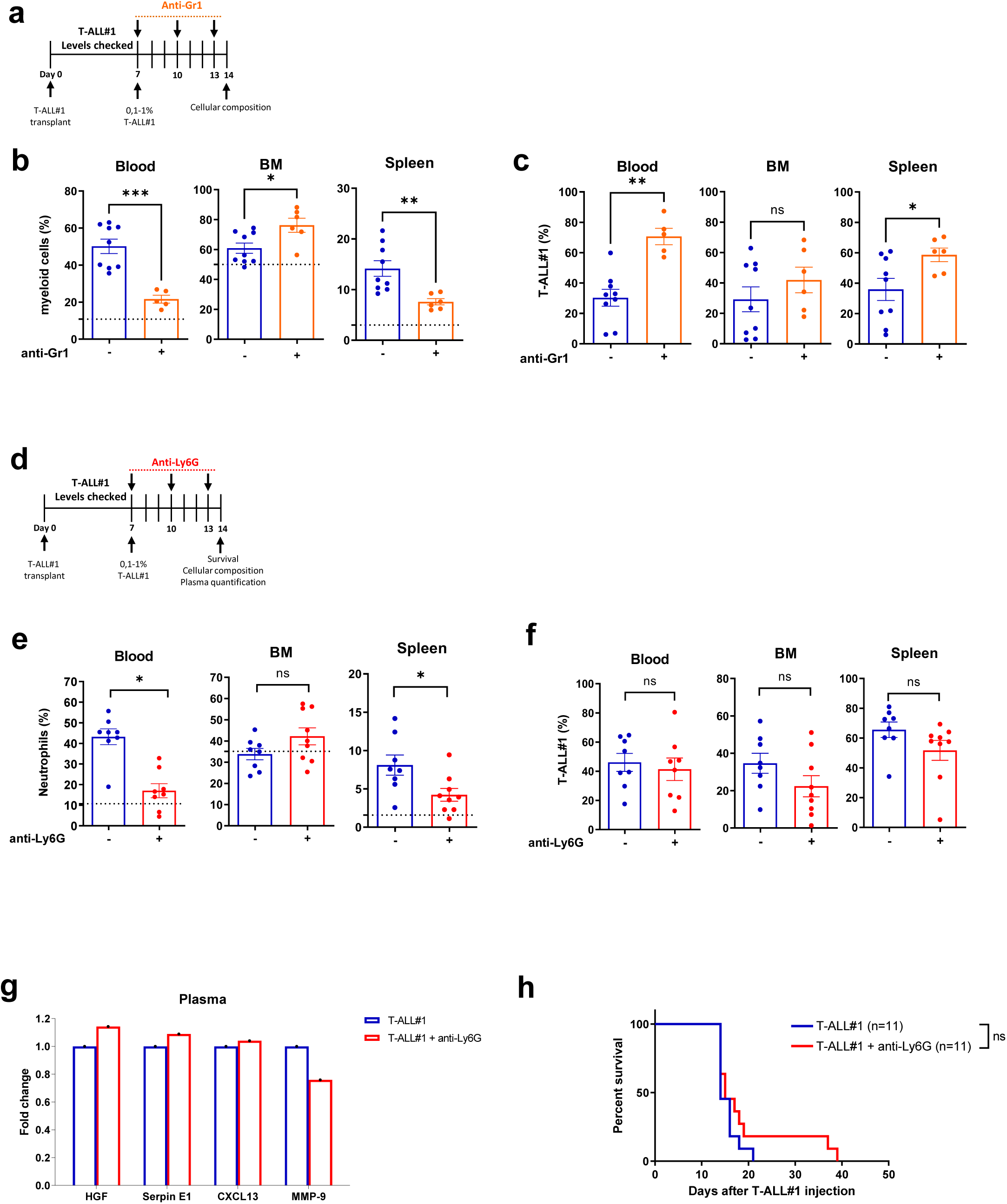
Neutrophil depletion does not impact T-ALL progression, plasma molecule secretion and mice survival. (a) Experimental design in which mice were treated for one week with 3 intraperitoneal injections of 100µg of anti-Gr1 depleting or isotype control antibodies between day 7 and day 13 after T-ALL#1 injection and were sacrificed at day 14. (b-c) Percentage of myeloid cells (CD11b^+^) (b) and of T-ALL#1 cells (c) in the blood, BM and spleen from anti-Gr1 (n=6) or isotype-treated (n=9) mice. The dotted line represents the percentage of cells in non-leukemic control mice. Shown are mean ± SEM from 2 independent experiments. (d) Similar protocol performed with anti-Ly6G or isotype antibodies. (e-f) Percentage of neutrophils (e) and T-ALL#1 cells (f) in blood, BM and spleen from anti-Ly6G (n=8) or isotype-treated (n=9) mice in 2 experiment. Shown are mean ± SEM from 2 independent experiments. g) Measurement of HGF, Serpine E1, CXCL13 and MMP-9 in the plasma from anti-Ly6G and isotype-treated mice identified by cytokine array. Each dot represents one cytokine array experiment. 2 plasma samples were pooled for each condition. (h) Kaplan-Meier survival curves of T-ALL#1 mice treated with anti-Ly6G or isotype. Here, an intensified neutrophils depletion was performed with a daily injection of 100µg anti-Ly6G (n=11 mice) or isotype (n=11 mice) antibodies for 5 days between day 7 and day 11 in 2 independent experiments, (log-rank Mantel-cox test was applied for statistical analysis). *p<0.05, **p<0.01, ***p<0.001 (two-tailed Mann-Whitney U test).

Unlike anti-Gr1, anti-Ly6G has the advantage to target neutrophils specifically ^33^. Anti-Ly6G has commonly been associated with a lower efficiency but higher specificity than anti-Gr1 to deplete neutrophils ^34^. When applying the same protocol as for anti-Gr1 treatment (Fig. 2d), a selective decrease (2-fold) of blood and spleen neutrophils was observed after anti-Ly6G treatment (Fig. 2e and Supplementary Fig. S2c-d). Consistently with anti-Gr1 results, neutrophils depletion did not impact T-ALL burden in the blood, BM and spleen (Fig. 2f and Supplementary Fig. S2c), neither modified the inflammatory molecules secreted in the plasma of anti-Ly6G-treated mice (Fig. 2g and Supplementary Fig. S2e).

Since neutrophils depletion was not complete after 3 injections of anti-Ly6G antibody, we evaluated the effect of a daily treatment with anti-Ly6G over 5 days from day 7 to day 11 on T-ALL burden and survival of mice. We found that a greater neutrophils depletion did not modify the survival of anti-Ly6G-treated mice that succumbed of T-ALL at similar time and with identical T-ALL burden compared to isotype-treated mice (Fig. 2h and data not shown). These results demonstrate that blood neutrophils are dispensable for T-ALL progression *in vivo* and suggest that other cell types such as resident monocytes and/or macrophages, likely contribute to T-ALL growth.

### Phagocytic cells support T-ALL without modulating molecular plasma secretion or mice survival

Based on our previous findings that the increase of blood resident monocytes was correlated with the T-ALL growth *in vivo*, we hypothesized that resident monocytes would contribute to T-ALL progression. Hence, we used clodronate liposomes that are known to deplete all phagocytic cells including resident monocytes and macrophages ^25^. The same protocol as anti-Ly6G treatment was applied and T-ALL burden was assessed after clodronate or vehicle treatment (Fig. 3a). In clodronate-treated T-ALL#1 mice, resident monocytes and macrophages significantly decreased without affecting inflammatory monocytes and neutrophils levels in blood, BM and spleen (Fig. 3b-c and Supplementary Fig. S3a-b). Macrophages and resident monocytes depletion strikingly diminished T-ALL burden in all tested cell compartment (Fig. 3d). Nevertheless, depleting specific phagocytic cells from mice with established T-ALL was not sufficient to decrease molecular plasma secretions or to impact mice survival (Fig. 3e-f and Supplementary Fig. S3c). These results demonstrate that leukemia associated resident monocytes and macrophages support established T-ALL progression *in vivo*. However, the phagocytic cell-mediated decreased T-ALL burden was not sufficient to decrease the molecular plasma microenvironment and the mice survival was unchanged, indicating they play significant but not major role in supporting T-ALL.

**Fig. 3:**
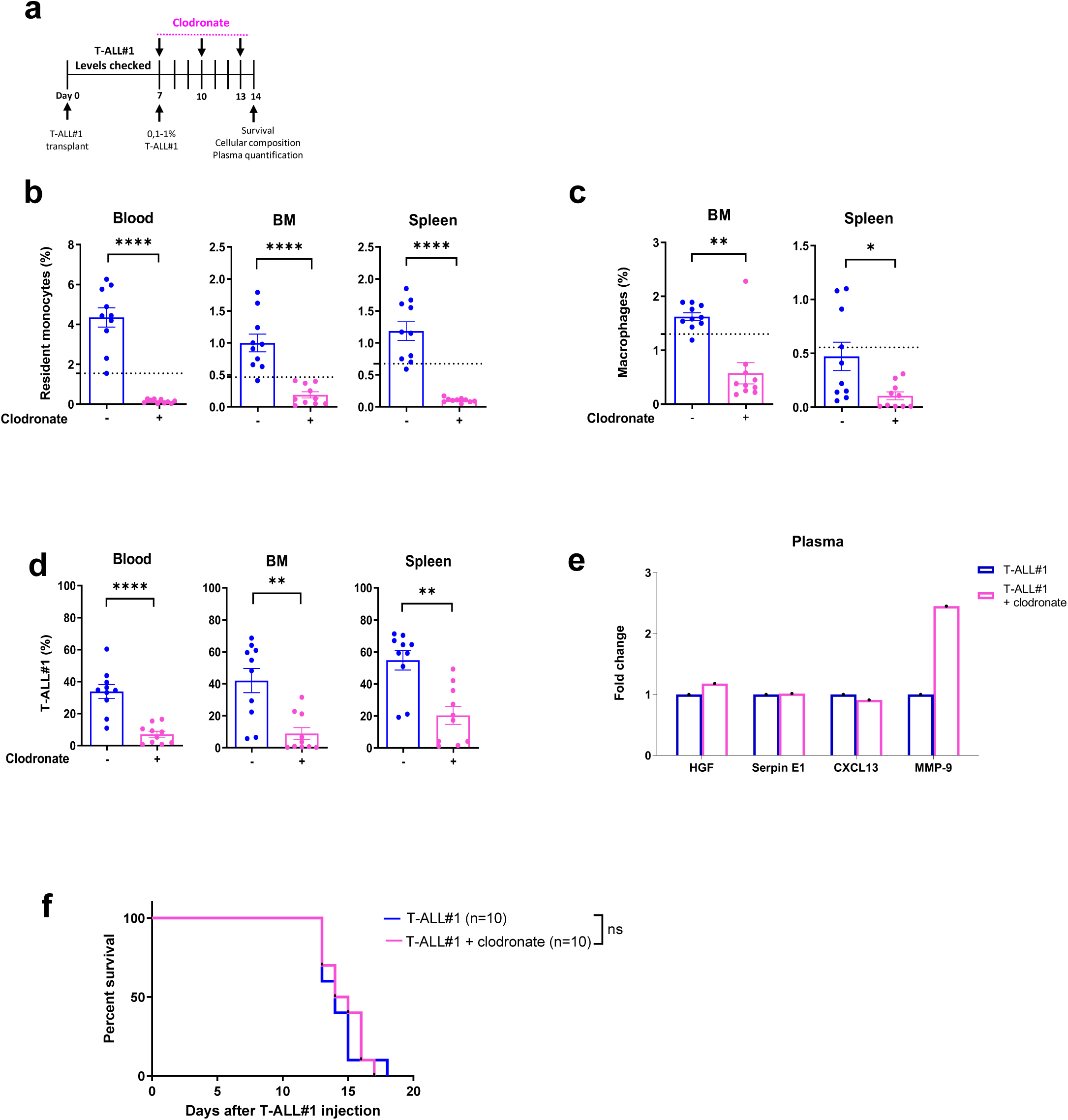
Phagocyte depletion with clodronate liposomes decreases T-ALL progression but does not impact molecule secretion in the plasma or mice survival. (a) Mice were treated for one week with 3 intraperitoneal injections of 1mg of clodronate liposomes or saline liposomes (vehicle) between day 7 and day 13 after T-ALL#1 injection and were sacrificed at day 14. (b-c) Percent of resident monocytes in the blood, BM and spleen (b) and percent of macrophages (Gr1^-^ CD115^-^ F4/80^+^ SSC^low^) in the BM and spleen (c) in clodronate (n=10) or vehicle-treated (n=10) mice in 2 experiments. The dotted line represents the percentage of cells in non-leukemic control mice. (d) Percent of T-ALL#1 cells in clodronate or vehicle-treated mice. Same mice as in (b-c). (e) Measurement of plasma HGF, Serpine E1, CXCL13 and MMP-9 levels from clodronate or vehicle-treated mice identified by cytokine array. Each dot represents one cytokine array experiment. 3 plasma samples were pooled for each condition. (f) Kaplan-Meier survival curves for T-ALL mice treated with clodronate or vehicle (10 mice in each treatment group in 2 independent experiments, log-rank Mantel-cox test was applied for statistical analysis). All data represent means ± SEM. *p<0.05, **p<0.01, ****p<0.0001 (two-tailed Mann-Whitney U test).

### NECA targets the cellular microenvironment and prevents leukemia-associated inflammation

Experimental studies have shown that adenosine receptor (AR) agonists are promising pharmacological modulators of disease-associated inflammation and immune responses ^35^. Adenosine is known to mediate anti-inflammatory effects via the adenosine A2a receptor (A2aR) ^36^. Our results show that mice injected with T-ALL#1 and T-ALL#2 significantly increase blood neutrophils, resident monocytes and inflammatory molecules. However, targeted depletion of neutrophils or resident monocytes does not significantly impact inflammatory molecule levels or confer a survival benefit to the leukemic mice. To focus more on the pro-leukemic effect of inflammatory molecule secretion in T-ALL, we treated T-ALL-transplanted mice with 5’-(N-ethyl-carboxamido)adenosine (NECA), a non-selective adenosine receptor agonist, known to decrease inflammatory factors secretion ^37–39^.

To exclude a direct interaction between NECA and leukemic cells, we first treated T-ALL#1 or T-ALL#2 cells *in vitro* with increasing NECA doses. T-ALL cells were counted by FACS analysis at different time points. Overall, similar leukemic cell growth was observed with or without NECA treatment suggesting that NECA does not interfere directly on T-ALL growth (Fig. 4a and Supplementary Fig. 4a). These results could be explained by the absence or low of expression of A2aR in T-ALL cells recovered from transplanted mice compared to BM and blood cells, in particular compared to myeloid cells (Fig. 4b and Supplementary Fig. S4b), suggesting leukemic cells are not sensitive to NECA. Next, we treated T-ALL-transplanted mice with NECA using a similar injection protocol as previously used with antibody experiments (Fig. 4c).

**Fig. 4:**
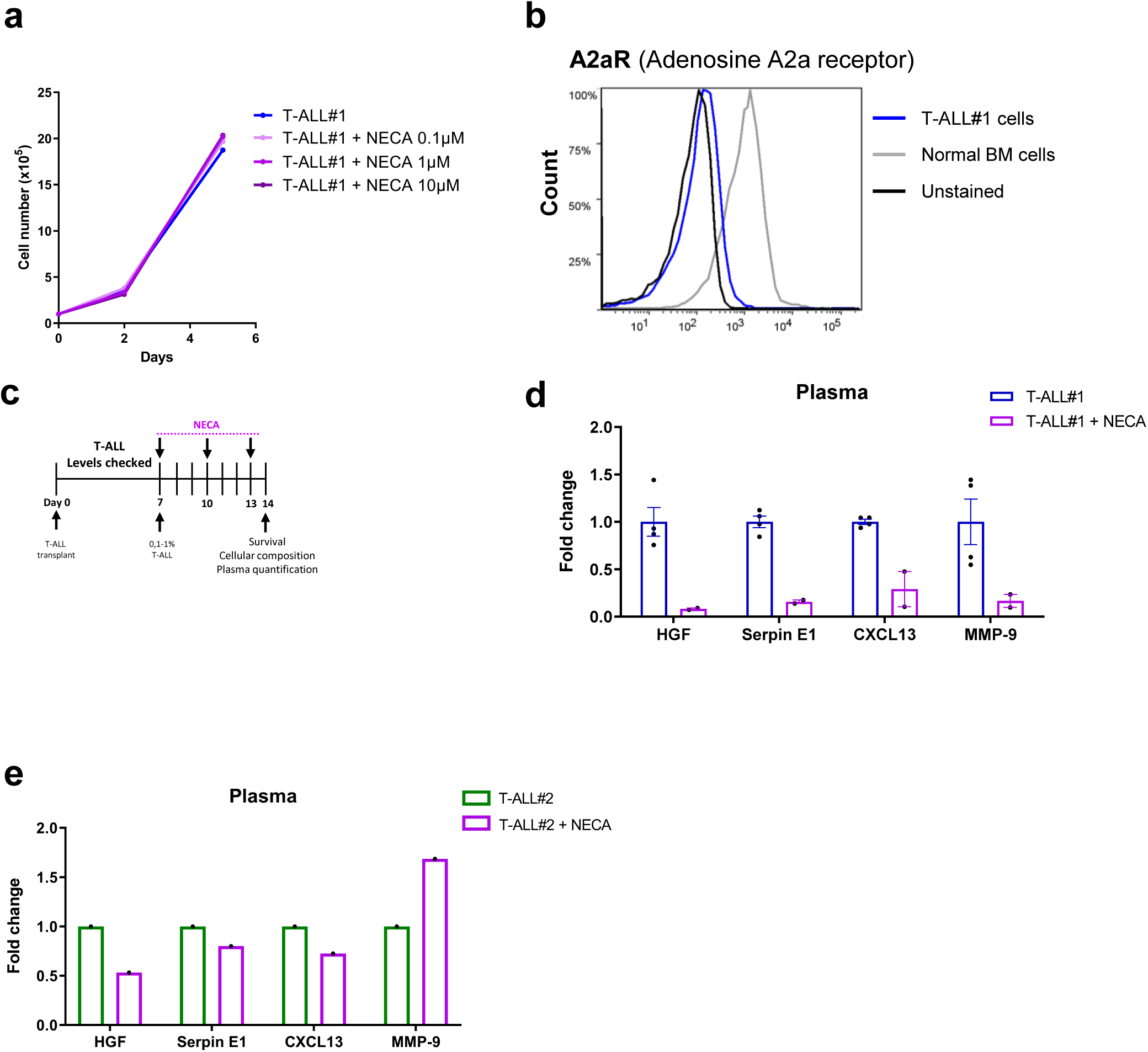
NECA acts indirectly on T-ALL cells and decreases molecule secretion in the plasma. (a) Growth curve of T-ALL cells cultured on MS5 stromal cells in the presence (TALL#1+NECA) or absence (T-ALL#1) of 10µM, 1µM or 0.1µM of NECA each time point is the median of triplicates. (b) Cell surface expression of Adenosine A2a receptor (A2aR) on T-ALL#1 cells and on normal mouse BM cells compared to unstained negative control. (c) Experimental design in which mice were treated for one week with 3 intraperitoneal injections of NECA or vehicle between day 7 and day 13 after T-ALL#1 injection and were sacrificed at day 14. (d) Measurement of HGF, Serpine E1, CXCL13 and MMP-9 levels in the plasma from NECA or vehicle-treated mice after T-ALL#1 injection. Each dot represents one cytokine array experiment and 3 or 4 plasma samples were pooled for each condition. All data represent means ± SEM. (e) same as in (d) in the plasma from NECA or vehicle-treated mice after T-ALL#2 injection. 2 plasma samples were pooled for each condition.

NECA-treated mice exhibited a strong reduction of the inflammatory molecular response observed in T-ALL models (Supplementary Fig. S4c), as expected from the fact that A2aR stimulation inhibits the transcriptional activity of NF-κB consequently suppressing the expression of pro-inflammatory cytokines ^36^. In particular, HGF was significantly decreased (12.3-fold, most decreased molecule) as were Serpin E1 (6.3-fold), CXCL13 (3.5-fold) and MMP-9 (6-fold) (Fig. 4d). In NECA-treated mice after T-ALL#2 injection, we also observed a decrease in several inflammatory molecules, though it was much more limited, with a 2-fold decrease of HGF for instance, but no decrease of Serpin E1, CXCL13 and MMP-9 (Fig. 4e and Supplementary Fig. 4d). On the cellular level, NECA treatment induced a decrease of blood percentage of neutrophils and a slight, albeit not significant, decrease of blood monocytes (Supplementary Fig. 4e).

Collectively these data show that NECA slightly modifies the cellular microenvironment of T-ALL and strongly inhibits the molecular inflammatory environment induced by the leukemia in *vivo*.

### Effect of NECA on T-ALL progression

To understand the impact of NECA on T-ALL growth *in vivo*, we analyzed T-ALL cell levels and viability in the blood, BM and spleen of T-ALL transplanted mice. NECA treatment of T-ALL#1 mice significantly diminished the weight of spleen associated with a decrease of T-ALL#1 burden in blood, BM and spleen at day 14 (Fig. 5a-c). Furthermore, NECA treatment (4 injections between day 7 and day 15) conferred a low but significant survival benefit, as NECA-treated mice succumbed from this aggressive leukemia 3 to 5 days after (day 18) compared to vehicle injected control mouse (Fig. 5d). The recovered mice had leukemic cell infiltration in their BM, Blood and spleen (Supplementary Fig. S5a). Interestingly, transplantation of BM T-ALL cells from NECA-treated T-ALL or control mice showed that all secondary recipient mice died at the same time, regardless of the leukemic cell origin, on average after 14 days (Supplementary Fig. S5b), suggesting NECA treatment did not induce specific intrinsic modifications on T-ALL cells but rather a transient response of leukemic cells to the NECA-mediated microenvironment changes.

**Fig. 5:**
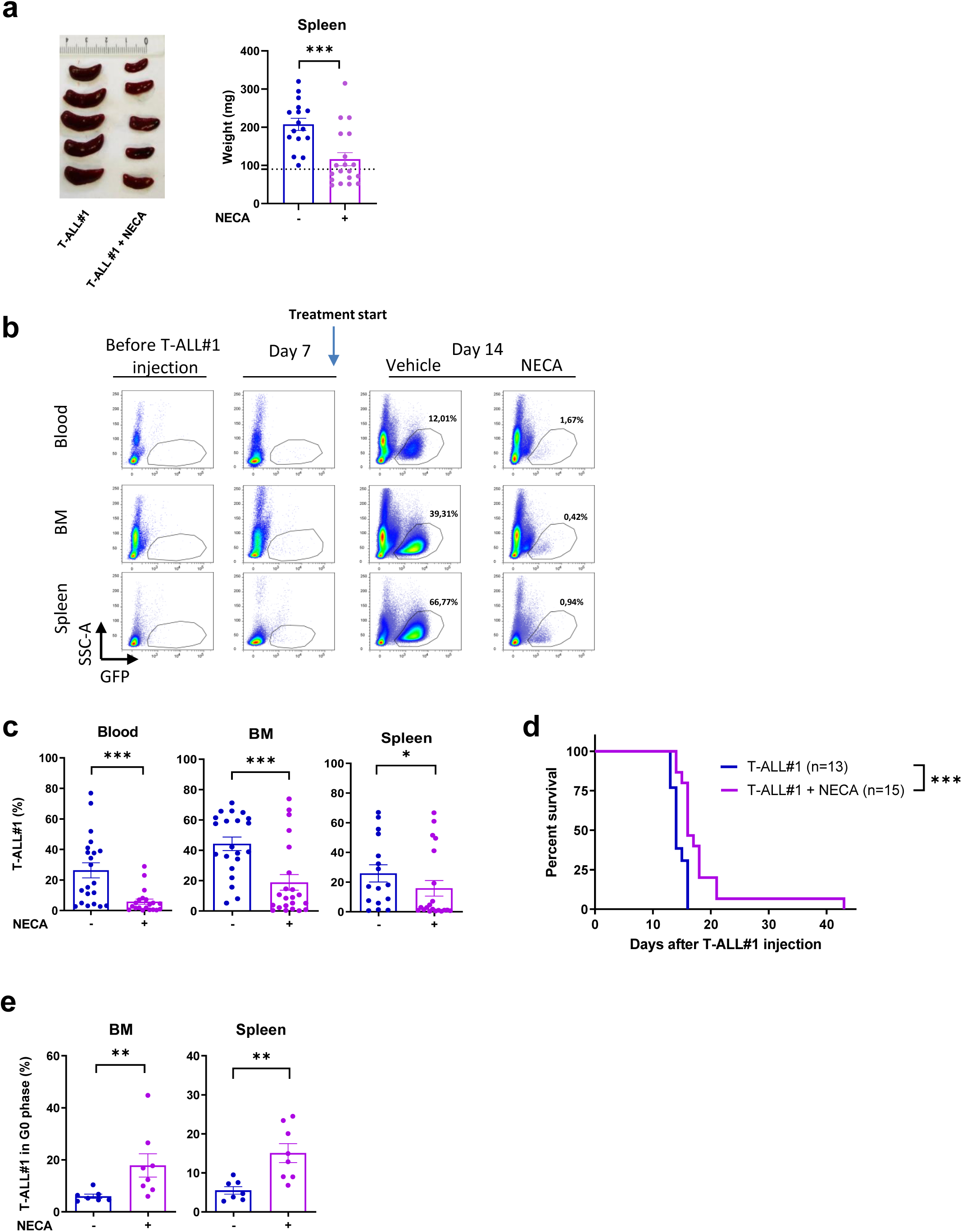
NECA decreases T-ALL progression and increases leukemic mouse survival. (a) Spleen weight and representative sizes in NECA (n=19) and vehicle-treated (n=16) mice from 4 independent experiments. (b) Representative flow cytometry plots before, and 7 and 14 days after T-ALL#1 injection in the blood, BM and spleen of NECA and vehicle-treated mice. (c) Percent of T-ALL#1 cells in the blood, BM and spleen of NECA (n=22) and vehicle-treated (n=20) mice in 6 independent experiments for blood and BM. (d) Kaplan-Meier survival curve of NECA (n=15) and vehicle-treated (n=13) mice after 4 intraperitoneal injections of NECA between day 7 and day 15 in 3 independent experiments. Log-rank Mantel-Cox statistical analysis was applied. (e) Percent of T-ALL#1 cells in G0 phase by flow cytometry in NECA (n=8) and vehicle-treated (n=7) mice in 2 independent experiments. All data represent means ± SEM. *p<0.05, **p<0.01, ***p<0.001 (two-tailed Mann-Whitney U test).

To validate that the impact of NECA treatment was not specific to the NOTCH1-driven T-ALL#1 model, mice injected with T-ALL#2 were also treated with NECA, starting when the first T-ALL#2 cells were detected in the blood until sacrifice (day 36). In this model NECA only induced a slight reduction of T-ALL burden that was only significant in the BM compartment (Supplementary Fig. S5c), maybe due to the weaker effect of NECA in the inflammatory molecule production in this T-ALL#2 model (Fig. 4e and Supplementary Fig. S4d).

We next investigated the proliferation and apoptosis of leukemic cells after NECA treatment in BM and spleen. In both organs, T-ALL#1 cells were less activated in NECA-treated mice, with higher leukemic cells in the G0 cell cycle phase and conversely lower proliferating Ki-67^+^ cells (Fig. 5e and Supplementary Fig. S5d-e). Besides, higher levels of apoptotic Annexin V^+^ leukemic cells were measured although only significant in the spleen (Supplementary Fig. S5f).

Altogether, these findings demonstrate that targeting the inflammatory microenvironment triggers a decrease of T-ALL cell burden mainly by reducing their proliferation and blocking them in G0 phase allowing a survival benefit of mice with leukemia.

### NECA decreases inflammatory pathways and identifies HGF as a potential molecular actor in T-ALL

To gain insight into the molecular regulation of NECA dependent inflammation in T-ALL, we investigated gene expression in T-ALL cells isolated from BM of NECA or vehicle-injected mice on day 14, the day after the last NECA injection, using microarray gene expression profiling. Principal component analysis (PCA) showed that NECA treatment induced transcriptional differences between samples. In the PCA score plot, the two groups were well separated with principal component (PC) 1 reaching 84.4%indicating that NECA treatment induces a significant alteration of various gene panels in T-ALL cells (Fig. 6a). Segregation of samples is validated by hierarchical clustering (Fig. 6b). Gene set enrichment analysis (GSEA) that upregulated DEGs (differential expressed genes) in vehicle compared to NECA injected-mice were significantly associated with KRAS signaling, type I (interferon alpha), and type II (interferon gamma) responses in vehicle-treated mice (Fig. 6c). As KRAS is a known pro-inflammatory tumor microenvironment modulator and interferon responses are well identified in several inflammatory pathways ^40–42^, the transcriptional difference in the inflammatory response in NECA-treated and untreated T-ALL#1 cells was consistent with the reduced inflammatory factor environment and T-ALL progression observed in NECA-treated mice. Over-representation using ingenuity pathway analysis (IPA) highlighted 61 pathways of interest with significant Fisher-exact adjusted p-value. Among those whose activation status could be calculated using z-score algorithm and several ones were associated with inflammatory response and cell cycle (Supplementary Fig. S6). Interestingly, pathway analysis showed that HGF signaling was among the top pathways preferentially inhibited in T-ALL#1 from NECA-treated mice (Supplementary Fig. S6), in line with the major decrease of HGF in the plasma of NECA-treated mice (Fig. 4d). Consistent with this, quantification analysis of HGF in the mouse plasma during T-ALL progression showed that the HGF was increased only 14 days after T-ALL#1 injection by 68-fold with T-ALL disease and decreased by 22-fold after NECA treatment (Fig. 6d).

**Fig. 6:**
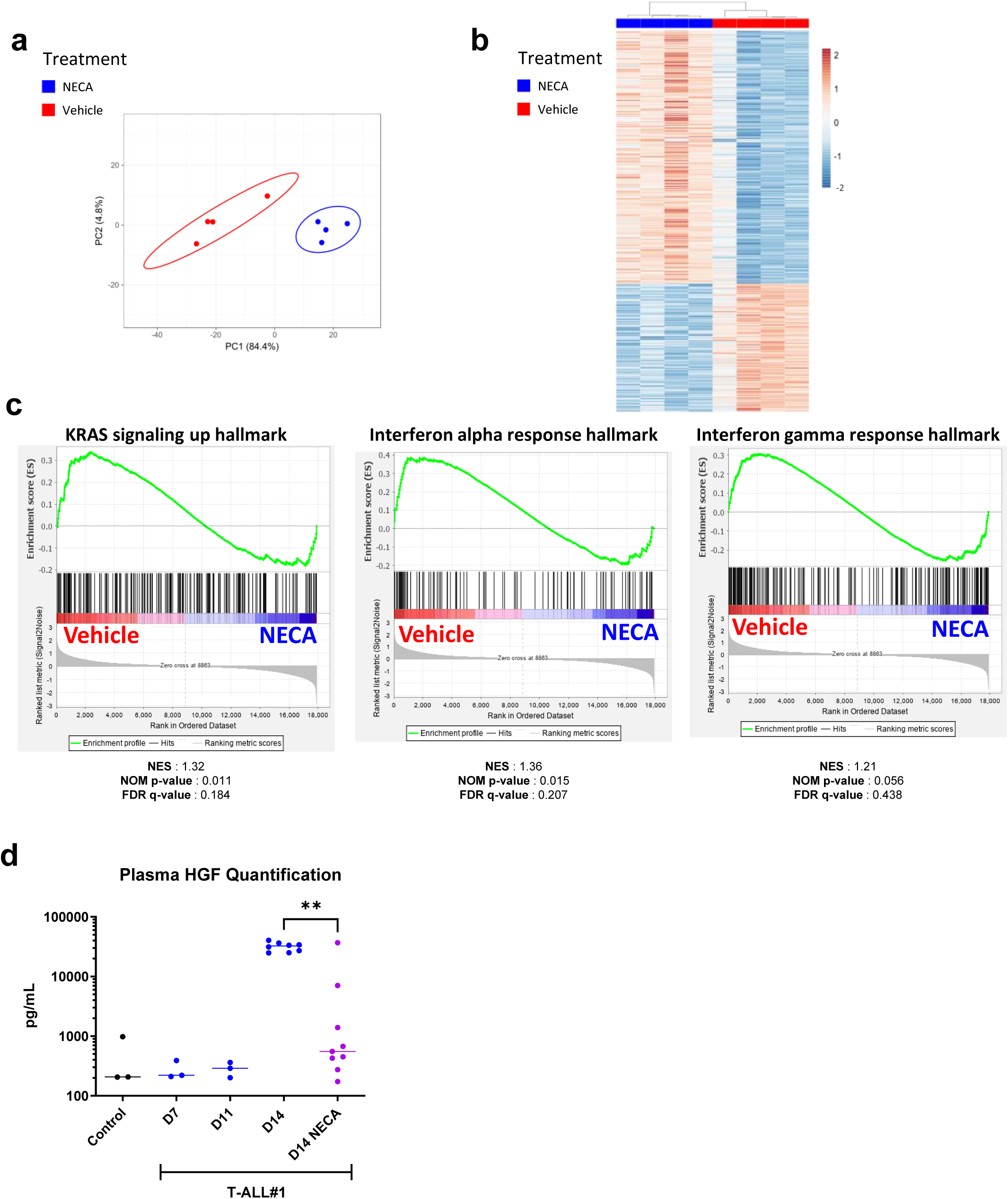
NECA treatment decreases inflammatory pathways at the transcripomic level and identifies HGF as a potential molecular actor in T-ALL. (a) Principal component analysis (PCA) based on Pearson’s correlation of the transcriptomic profile of T-ALL#1 cells from NECA (blue) or vehicle-treated (red) mice. Clusters of quadruplicate values are represented in circles. (b) Heatmap of the genes expressed with a two-fold difference between NECA or vehicle condition. (c) Gene Set Enrichment Analysis (GSEA) positively enriched in cells treated with vehicle compared to cells from NECA treatment. The normalized enrichment score (NES), nominal p-value (NOM p-value) and False Discovery Rates q-value (FDR q-value) are indicated for each GSEA plot. (d) Quantification of plasma HGF in non-leukemic (Control, n=3) mice and in T-ALL#1 leukemic mice 7, 11 and 14 days post T-ALL#1 injection in vehicle- (n=8) or NECA-treated (n=9) mice from 3 independent experiments. All data represent means ± SEM. **p<0.01 (two-tailed Mann-Whitney U test).

Collectively, these studies indicated that diminishing the inflammatory microenvironment with NECA treatment in T-ALL mice regulates several pathways associated with cancer induced inflammation including HGF/cMet signaling, interrogating the role of this pathway in T-ALL progression.

### HGF/cMet signaling pathway induces survival and proliferation of T-ALL#1 cells

We next tested whether the HGF signaling pathway could act as a major activator of T-ALL cells. Flow cytometry analysis and western blot showed that T-ALL#1 cells express cMet, the HGF receptor, indicating that this growth factor could interact with T-ALL#1 cells (Fig. 7a and Supplementary Fig. S7a). It is described that the binding of HGF to c-MET induces the dimerization of c-MET that enables its intracellular kinase domains (KDs) to undergo autophosphorylation leading to activation of downstream signaling pathways ^43^. We next determined the levels of c-Met phosphorylation in T-ALL#1 cells *in vitro* in coculture with MS5 stromal cells without external HGF addition as HGF is highly produced in MS5 cell supernatant (Supplementary Fig. S7b). We observed using flow cytometry analysis with a specific antibody to phospho-cMet that the phosphorylation of cMet was present in T-ALL#1 cells cocultured with MS5 stromal cells (Fig. 7b) suggesting that HGF could act on leukemic cell growth in the context of microenvironment stimulation. To test this hypothesis, we used the c-Met inhibitor SU11274 to block cMet phosphorylation and function in T-ALL#1 cells ^44^. After treatment with SU11274 for 24h, T-ALL cells had decreased cMet phosphorylation associated with a significant decrease of T-ALL#1 growth (Fig. 7c-d). Interestingly SU11274 exerted its action on leukemic cell growth through cell death as SU11274 treatment doubled apoptosis levels (Fig. 7e) but did not modify the leukemic cell cycle progression except for an increased proportion of cells in sub-G1 phase when compared with the vehicle control condition (Fig. 7e, and Supplementary Fig. S7c).

**Fig. 7:**
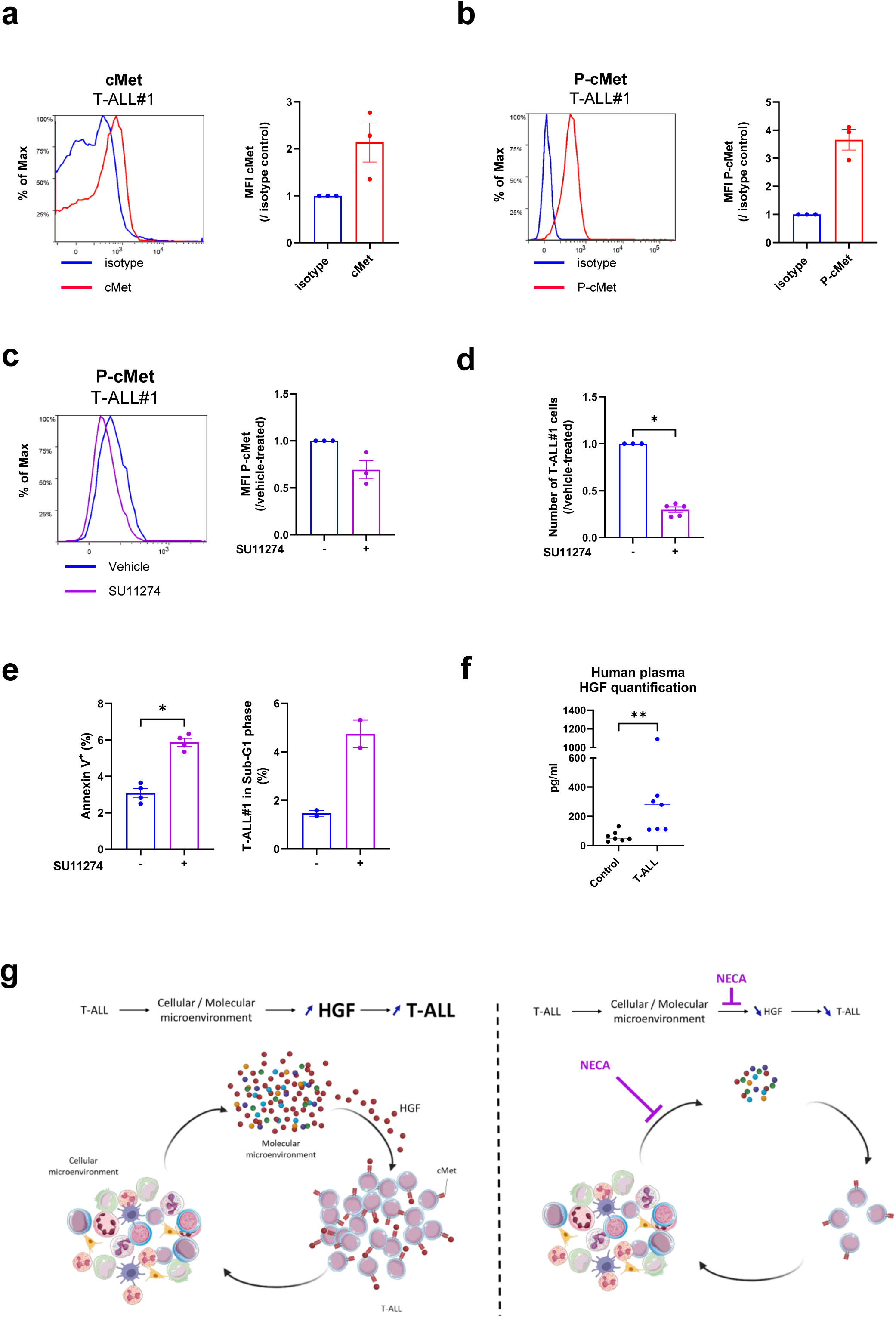
HGF/cMet signaling pathway, a new putative driver of T-ALL. (a) Cell surface expression of cMet on T-ALL#1 cells compared to isotype control by flow cytometry (left panel). Mean Fluorescence Intensity (MFI) of cMet expression on T-ALL#1 cells normalized by the MFI of the isotype control (right panel) in 3 independent experiments. (b) Same as in (a) for cMet phosphorylation (P-cMet) in T-ALL#1 cells co-cultured with MS5 cells compared to isotype control (left panel). MFI of P-cMet on T-ALL#1 cells co-cultured with MS5 cells normalized by the MFI of the isotype control (right panel) in 3 independent experiments. (c) Same as in (a-b) for P-cMet detection in presence or not (vehicle-treated T-ALL#1 cells) of 5µM SU11274 inhibitor during 24h co-cultured with MS5. MFI of P-cMet in SU11274 treated T-ALL#1 cells normalized to P-cMet MFI of T-ALL#1 vehicle-treated cells (right panel) in 3 independent experiments. (d) Number of T-ALL#1 cells with or without (vehicle) 5µM of SU11274 during 24h co-cultured with MS5. Representation was normalized on the number of T-ALL#1 vehicle-treated cells in 3 independent experiments. (e) Proportion of Annexin V positive T-ALL#1 cells in presence or not (vehicle-treated T-ALL#1 cells) of 5µM SU11274 inhibitor co-cultured with MS5 in 4 independent experiments (left panel). Proportion of T-ALL#1 cells in Sub-G1 phase of the cell cycle identified by flow cytometry in SU11274 or vehicle-treated cells in 2 independent experiments (right panel). (f) Quantification of HGF in the plasma of 7 control and 7 T-ALL human samples by multiplex analysis. (g) Overview of the role of inflammatory microenvironment on T-ALL progression on the basis of the results shown in this study. T-ALL induces a cellular and molecular inflammatory response from the microenvironment favoring the secretion of HGF which in turn supports T-ALL progression. NECA treatment inhibits inflammatory molecule secretion by the cellular microenvironment which causes an impairment of HGF release then unable to support T-ALL progression. Data a to e represent means ± SEM. Data f represent values and median. *p<0.05, **p<0.01 (two-tailed Mann-Whitney U test).

To evaluate the relevance of our observations in human T-ALL, cytokine array analysis was conducted to measure the circulating HGF levels in plasma of T-ALL patients in comparison with healthy donors. Interestingly, HGF was detected among the 20 highest increased inflammatory molecules in plasma of T-ALL patients compared to controls, (Supplementary Fig. S7d). A precise quantification of HGF in the plasma of 7 newly diagnosed T-ALL patients by Luminex assay showed that HGF was significantly increased (6-fold) compared to plasma controls (Fig. 7f).

Taken together, these results indicate that inhibiting the activation of HGF/cMet axis by SU11274 treatment reduces the viability of T-ALL#1 cells and reveals HGF/cMet signaling pathway as a putative driver of T-ALL expansion (Fig. 7g).

## DISCUSSION

In this issue, we show using T-ALL mouse models that leukemic progression is accompanied with a cellular and molecular inflammatory microenvironment that participate in leukemic cell expansion, thus favoring high leukemic burden.

T-ALL diversity driven by multiple oncogenic events can be at the origin of a variability in the response to drugs targeting the tumor cells ^8,10,45,46^. However, leukemia targeting drugs like chemotherapeutic compounds, albeit efficient in many patients, have been shown to induce negative side effects ^4^ and to trigger chemoresistance resulting in relapse with poor outcome ^47^. Thus, alternative therapeutic approaches have yet to be developed to improve patient treatments, supporting the importance of continuing to study T-ALL biology, and in particular leukemic cell interactions with their microenvironment to find therapeutically targetable original pathways.

In this context, inflammation is a common feature of many solid and hematological malignancies and is widely recognized as an important factor in tumor development, progression, metastasis, and resistance to treatment, *i.e.* through its ability to suppress antitumor immune responses ^48,49^. By contrast, inflammation related to leukemia-infiltrated tissues is not widely recognized as a feature of T-ALL. However, multiple studies show that an inflammatory microenvironment promotes leukemia progression. Indeed, genetically predisposed Pax5 haplo-insufficient mice can develop B-ALL when exposed to inflammatory environments ^50^. Also, statistical analysis and etiological studies have shown that enhanced inflammatory reaction in early childhood could induce wakening of ALL cells triggering leukemic development ^51^. Furthermore, NF-κB pathway activation, a major hub in the regulation of inflammation, correlates with rapid childhood T-ALL evolution in PDX models ^52^ underlying the importance of studying the role of inflammation in T-ALL.

Our current results show that there is an increase of the cellular and molecular inflammatory microenvironment correlating with T-ALL progression, raising the question of the identity of the actors and their contribution to leukemic expansion. Previous works showed that thymic Dendritic Cells (DC) play a supportive role of T-ALL through the activation of IGF1R pathway of leukemic cells ^14^. Besides, other type of myeloid cells like macrophages and monocytes can also support T-ALL initiation and progression in various organs ^25^ through integrin-mediated cell adhesion and FAK/PYK2 signaling ^53^. Although our data confirm that phagocytes including resident monocytes support T-ALL progression, targeting these cells did not improve mice survival. We also independently studied neutrophils, a key cell population in inflammation, and showed that these cells are not implicated in T-ALL progression, albeit participating in other hematological malignancies like Chronic Lymphocytic Leukemia ^54^ or B lymphomas ^55^. Studying the role of neutrophils and resident monocytes in T-ALL was also motivated by their crucial and complex role in infections ^56^ or in cancers ^57^ and since they are the most increased cell populations in the periphery of our leukemic mice. However, show that contrary to our expectations, depleting specific myeloid cells such as neutrophils or phagocytes did not significantly impact inflammatory molecule secretion or improve T-ALL mice survival. These results comfort the importance of the cellular and the molecular actors in the survival of the mice and that the secretion seems to be enrolled by other cell populations than myeloid cells in the microenvironment. Nevertheless, it is most likely that several cell populations together are implicated in the immune responses in T-ALL, indicating that targeting one specific cell type may not be sufficient to efficiently module inflammation in T-ALL. Moreover, stromal cells have been shown to be modified and secrete supporting factors for ALL cells ^17,58^. Interestingly, it has been recently shown that inflammatory macrophages can educate other macrophages through IFNγ signaling in the microenvironment to phagocyte T-ALL cells and decrease the leukemic progression ^59^. However, when macrophages are depleted with clodronate treatment in our study, a decrease in leukemic progression is observed suggesting the existence of other mechanisms to support T-ALL progression.

Altogether, our results show the existence of mechanisms that promote paracrine interactions between the microenvironment and T-ALL cells. In this context and to investigate the role of inflammation in T-ALL, we tested whether targeting the inflammatory molecule secretion would modify leukemic cell infiltration. The assets of using NECA in this context are that instead of targeting one specific ligand receptor interaction, we can globally analyze the impact of the entire molecular inflammatory microenvironment on T-ALL and eventually identify new signaling pathways impacting T-ALL cells regardless of a specific cell type in the microenvironment. Indeed, many studies show independently that interaction between receptors and ligands such as DL1, IL-18, IL-7, IGF1 and CXCL12 sustain proliferation, survival or initiation of T-ALL ^12,15–17,60,61^. Indeed, treating T-ALL mice with NECA, a non-selective adenosine receptor agonist which does not act on T-ALL cells, significantly decreased T-ALL progression and prolonged mouse survival. NECA treatment induces a marked reduction in the levels of various inflammatory molecules.

A key revelation of this study is the identification of HGF/cMet signaling as a potential driver of T-ALL. The study demonstrates that HGF levels are elevated in plasma of T-ALL mice and patients, and gene expression analysis following NECA treatment reveals a notable inhibition of the HGF/cMet signaling pathway in T-ALL cells. Moreover, using specific HGF/cMet signaling inhibitor, we reveal that this pathway is required for T-ALL growth, emphasizing the importance of molecule secretion in T-ALL microenvironment. Indeed, the activation of the HGF/cMet pathway induces cell survival and proliferation via tyrosine kinase cascades such as MAPK, PI3K/AKT or JAK/STAT ^62^. On top of its supportive role in hepatic cancers ^63^ or in the metastasis and growth process of solid tumors ^64^, the HGF/cMet signaling axis has been shown to also contribute to the progression of different types of leukemia like Acute Myeloid Leukemia ^65^, Chronic Lymphocytic Leukemia ^66^ and Adult T cell Leukemia ^67^. In addition, even if several secreted factors have been identified to play important role in the initiation or the progression of T-ALL, we believe that it is the combination of them that constitute a favoring soil for T-ALL expansion and targeting their action with combined inhibition of several signaling pathways at once could be a promising therapeutic approach in the field of T-ALL treatment.

Recent studies have shown that T-ALL cells contribute to the inflammatory activation of the microenvironment, namely via calcium signalization with the expression of STIM gene family ^24^. Deletion of STIM in leukemic cells decrease inflammatory response inducing the prolongation of mice survival. This strategy is complementary to our study with NECA. Both studies rely on decreasing inflammatory microenvironment of T-ALL except that we target inflammatory molecular secretion by cellular microenvironment when they target initiating inflammatory inducing event intrinsic to T-ALL. In both scenarios, decreasing inflammatory microenvironment in T-ALL decreases the leukemic burden and improves mice survival.

To sum up, we showed here that secreted inflammatory molecules supports T-ALL progression. We identified HGF/cMet signaling, as a potential pathway supportive of T-ALL progression. Our results support dual therapeutic approaches that could complement existing treatment protocols, either targeting the entire molecular secretion by the microenvironment or specifically inhibiting HGF/cMet signaling pathway.

## Contributors

CLM, LF, LR and VP performed experiments. CLM, LF, BU and VP analyzed data. CLM and MLG analyzed microarrays experiments. VB, CD and CD assisted with animal experiments. JC gave valuable advices during experiments. BP and AP provided human T-ALL samples. BU, VP and FP conceived and mentored the project. CLM, VP and FP wrote the manuscript. All authors critically reviewed the manuscript.

## Declaration of interest

The authors declare that they have no conflict of interest.

## Acknowledgements

We are grateful to the staff of the iRCM animal platform and to N. Dechamps and J. Baijer of the iRCM cytometry and cell sorting platform, for their help in experiments. The authors thank J. Ghysdael and C. Tran Quang for providing ICN1/T-ALL#1 cells and C. Smith and V. Asnafi for LMO1tg/ SIL-TAL1tg mouse models. We acknowledge Dr Peter Aplan for giving his authorization to use the TAL1-LMO1 mice in our project. The authors would like also to thank the human donors for allowing the use of their plasma in our research. Microarray experiments were greatly facilitated by D. Lewandowski.

## Data sharing statement

Data collected for these studies will be made available upon request to the corresponding author.

## SUPPLEMENTARY FIG. LEGENDS

**<u>Supplementary Fig. S1:</u>** (a) Kaplan-Meier survival curve of T-ALL#1 injected mice (n=36). 50% of mice deceased at day 14 (dotted line). (b) Gating strategy performed by flow cytometry to identify T cells, B cells, neutrophils, inflammatory monocytes and resident monocytes. (c) Correlation between the percent of either blood inflammatory monocytes, B cells and T cells and the percent of blood T-ALL#1 cells. Results from 60 mice are shown (Spearman’s r correlation test was applied). (d) Absolute number of neutrophils and resident monocytes per µl of blood at day 14. Shown are the mean ± SEM of 10 control and 24 T-ALL#1 mice from 8 independent experiments. (e) Typical flow cytometry analysis of T-ALL#2 cells from BM T-ALL#2 injected mice. (f) Percent of T-ALL#2 cells in the blood, BM and spleen of T-ALL#2 injected mice at sacrifice (4 or 5 mice). (g) Top 20 most increased inflammatory molecules in T-ALL#1 mice plasma measured by cytokine array. 2 plasma samples were pooled for control in 2 independent experiments. 3 or 4 plasma were pooled for T-ALL#1 in 4 independent experiments. The fold increase of each molecule in leukemic mice compared to controls is represented in grey dotted line. (h) Top 20 most increased inflammatory molecules in T-ALL#2 mice plasma measured by cytokine array. Fold increase of each molecule in leukemic mice compared to controls is represented in grey dotted line. 2 plasma samples were pooled for controls in 2 independent experiments and 2 plasma samples were pooled for T-ALL#2 in 1 experiment. All data represent means ± SEM. ****p<0.001 (two-tailed Mann-Whitney U test).

**<u>Supplementary Fig. S2:</u>** (a) Absolute number of myeloid (left panels) and T-ALL#1 (right panels) cells per µl of blood, in the BM of two femurs, or in the spleen (9 control mice and 6 anti-Gr1 treated mice in 2 independent experiments). (b) Percent of neutrophils (left), inflammatory monocytes and resident monocytes (right) in anti-Gr1 or isotype-treated mice in blood, BM and spleen (9 controls mice and 6 anti-Gr1 treated mice in 2 independent experiments). (c) Absolute numbers of neutrophils (left) and T-ALL#1 (right) cells per µl of blood, in the BM of two femurs, or in the spleen (8 controls mice and 8 anti-Ly6G treated mice in 2 independent experiments). (d) Percent of resident monocytes and inflammatory monocytes in anti-Ly6G or isotype-treated mice in blood, BM and spleen (6 control and 5 anti-Ly6G treated mice in 1 experiment). (e) Top 20 most decreased inflammatory molecules measured by cytokine array in the plasma of anti-Ly6G or isotype-treated (T-ALL#1) mice. Fold decrease of each molecule is represented in grey dotted line. Every dot represents one cytokine array experiment. 2 plasma samples were pooled for each condition in 1 experiment. All data represent means ± SEM.*p<0.05, **p<0.01, ***p<0.001 (two-tailed Mann-Whitney U test).

**<u>Supplementary Fig. S3:</u>** (a) Absolute numbers of resident monocytes, and T-ALL#1 cells per µl of blood, in the BM of two femurs, or in the spleen. Absolute numbers of macrophages in the BM and the spleen (10 mice in control and in clodronate treated group in 2 independent experiments). (b) Percent of neutrophils and inflammatory monocytes in the blood, BM and spleen (10 mice in each group in 2 independent experiments). (c) Top 20 most decreased inflammatory molecules measured by cytokine array in the plasma of clodronate or vehicle-treated mice. Fold decrease of each molecule is represented in grey dotted line. Each dot represents one independent cytokine array experiment. 3 plasma samples were pooled for each condition. All data represent means ± SEM.*p<0.05, **p<0.01, ***p<0.001, ****p<0.0001 (two-tailed Mann-Whitney U test).

**<u>Supplementary Fig. S4:</u>** (a) Growth curve of T-ALL#2 cells cultured on MS5 stromal cells in the presence (T-ALL#2 + NECA) or absence (T-ALL#2) of 10, 1 or 0,1 µM of NECA. (b) Mean Fluorescence Intensity (MFI) of Adenosine A2a receptor (A2aR) associated with blood and BM immune cells (inflammatory monocytes Ly6C^+^, resident monocytes Ly6C^+^, neutrophils, lymphoid cells) compared to T-ALL#1 cells. (c-d) Top 20 most decreased inflammatory molecules measured by cytokine array in the plasma of NECA compared to vehicle-treated mice after T-ALL#1 (c) or T-ALL#2 injection (d). Each dot represents one independent cytokine array experiment with 3 or 4 plasma (c) or 2 plasma pooled for each condition (d). Fold decrease of each molecule is represented in grey dotted line. (e) Percent of neutrophils, inflammatory monocytes and resident monocytes in NECA compared to vehicle-treated mice in blood after T-ALL#1 injection (7 control and 8 NECA-treated mice in 2 independent experiments). Data c and e represent means ± SEM. *p<0.05 (two-tailed Mann-Whitney U test).

**<u>Supplementary Fig. S5:</u>** a) Percent of T-ALL#1 cells 3 days after the last NECA injection (day 18) (related to Fig. 5d). Results are from 3 NECA-treated mice in 1 experiment. b) Experimental procedure of secondary transplantation of BM T-ALL#1 cells (upper panel) and Kaplan-Meier survival curve of secondary transplanted mice from NECA treated or vehicle BM T-ALL#1 cells (lower panel) (8 from vehicle and 6 from NECA-treated mice in 2 independent experiments). (c) Percent of T-ALL#2 cells in blood, BM and spleen of NECA (n=5) compared to vehicle-treated mice (n=5). (d) Representative flow cytometry plot for cell cycle analysis and (e) percent of proliferating Ki-67^+^ T-ALL#1 cells in NECA and vehicle-treated mice. Results are from 7 vehicle and 8 NECA-treated mice in 2 independent experiments. (f) Percent of apoptotic Annexin V^+^ T-ALL#1 cells in NECA and vehicle-treated mice. From 7 vehicle and 8 NECA-treated mice in 2 independent experiments. Data a and c represent values and median. Data e and f are represented with means ± SEM. *p<0.05, **p<0.01 (two-tailed Mann-Whitney U test).

**<u>Supplementary Fig. S6:</u>** (a) Pathways identified by IPA significantly decreased in NECA condition (bleu) or significantly increased in NECA condition (orange).

**<u>Supplementary Fig. S7:</u>** (a) Western blot showing cMet protein levels in T-ALL#1 cells in coculture with MS5. GAPDH was used as internal control. (b) HGF concentration in the supernatant of MS5 cell culture medium after 10 days of culture. (c) (left) Percent of proliferating Ki-67^+^ T-ALL#1 cells and (right) percent of T-ALL#1 cells in G0 phase in SU11274- and vehicle-treated cells. (d) Relative quantification of the top 20 most increased inflammatory molecules measured by cytokine array in the human T-ALL patient (n=4) or healthy donors (n=2).

**<u>Supplementary Table S1:</u>** Biological characteristics of human T-ALL samples used

**<u>Supplementary Table S2:</u>** List of PCR primers

**<u>Supplementary Table S3:</u>** List of antibodies used in flow cytometry

**<u>Supplementary Table S4:</u>** List of antibodies used in Simple Western

## Notes

### Competing Interest Statement

The authors have declared no competing interest.

