## Supplementary Tables for "T-cell acute lymphoblastic leukemia progression is supported by inflammatory molecules including Hepatocyte Growth factor"

### Supplementary Table S1

| T-ALL samples | Gender | Age | Genetic abnormalities |
|---|---|---|---|
| hT-ALL#1 | M | 14 | TLX3 |
| hT-ALL#2 | M | 10 | ND |
| hT-ALL#3 | F | 6 | ND |
| hT-ALL#4 | M | 5 | t(7;10) |
| hT-ALL#5 | M | 16 | TLX3 |
| hT-ALL#6 | M | 10 | NOTCH1 mutated |
| hT-ALL#7 | M | 16 | ND |

### Supplementary Table S2

| | Primers for TAL1 and LMO1 constructions | |
|---|---|---|
| | Forward | Reverse |
| TAL1 transgene | 5’-CACAGGGTCCTTGCCAGTC-3’ | 5’-CTACCCTGCAAACAGACCTC-3’ |
| LMO1 transgene | 5’-CGGGATCCGAGATGATGGTGCTGG-3’ | 5’-CCCAAGCTTACTGAACTTGGCATTCAAAGG-3’ |

### Supplementary Table S3

| Antigen | Clone | Source |
|---|---|---|
| B220-V500 | RA3-6B2 | BD Bioscience |
| CD115-PE | ASF98 | BioLegend |
| CD115-PE/Cyanine 7 | ASF98 | BioLegend |
| CD11b-BV605 | M1/70 | BioLegend |
| CD11b-PECyanine 7 | M1/70 | BioLegend |
| CD25-BV421 | PC61 | BioLegend |
| CD3-PerCP/Cyanine5.5 | 145-2C11 | BioLegend |
| CD44-APC/Cyanine 7 | IM7 | BioLegend |
| CD45.1-APC | A20 | BioLegend |
| CD45.2-FITC | 104 | BioLegend |
| CD4-BV421 | GK1.5 | BioLegend |
| CD4-PerCP/Cyanine 5.5 | GK1.5 | BioLegend |
| CD8a-APC | 53-6.7 | eBioscience |
| CD8a-PE | 53-6.7 | BD pharmingen |
| cMet-PE | eBioclone 7 | Invitrogen eBioscience |
| Phospho cMet-APC | MetY12341235-6F11 | Invitrogen |
| F4/80-AF700 | BM8 | BioLegend |
| Gr1-PerCP/Cyanine 5.5 | RB6-8C5 | BioLegend |
| Ly6C-APC | HK1.4 | Invitrogen |
| Ly6C-BV711 | HK1.4 | BioLegend |
| Ly6G-BV421 | 1A8 | BioLegend |

### Supplementary Table S4

| Antigen | Clone | Reference | Supplier |
|---|---|---|---|
| cMet | E4E4W | 82202S | Cell Signaling Technology |
| GAPDH | 14C10 | 2118 | Cell Signaling Technology |
