## Supplementary Figures for "T-cell acute lymphoblastic leukemia progression is supported by inflammatory molecules including Hepatocyte Growth factor"

#### Supplementary Fig. S1

**a**

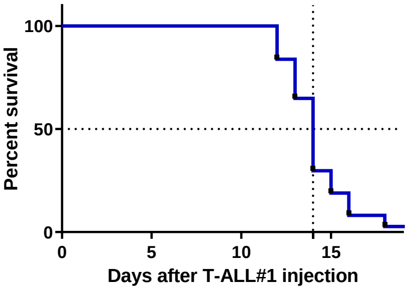


**b**

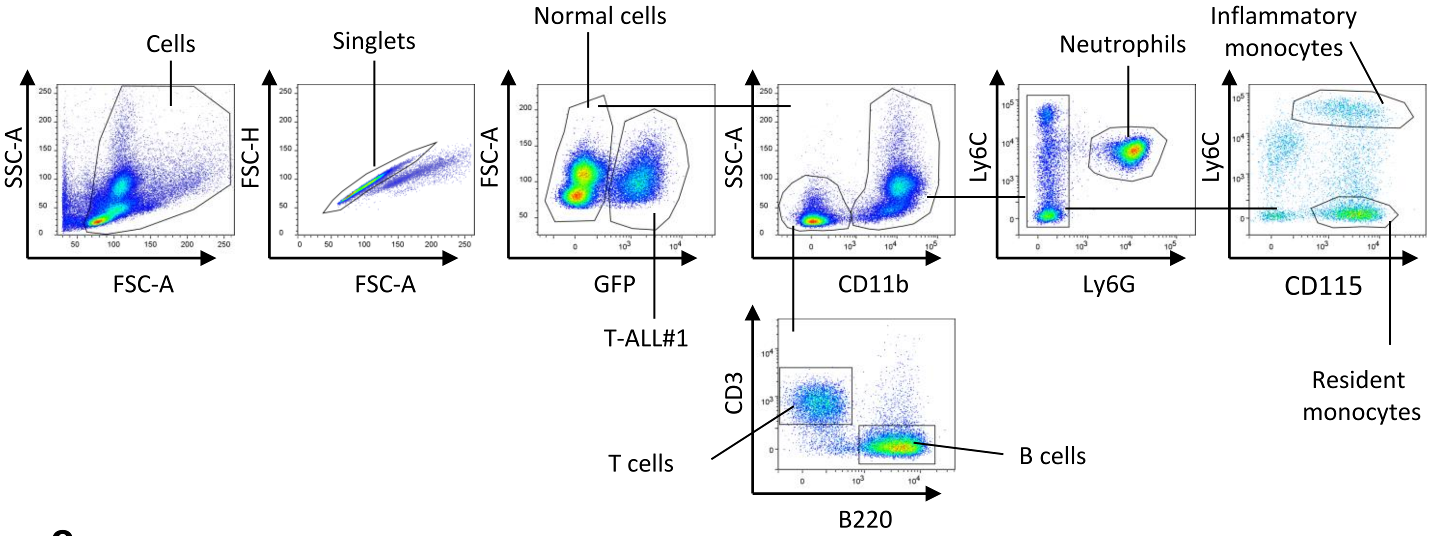


**c**

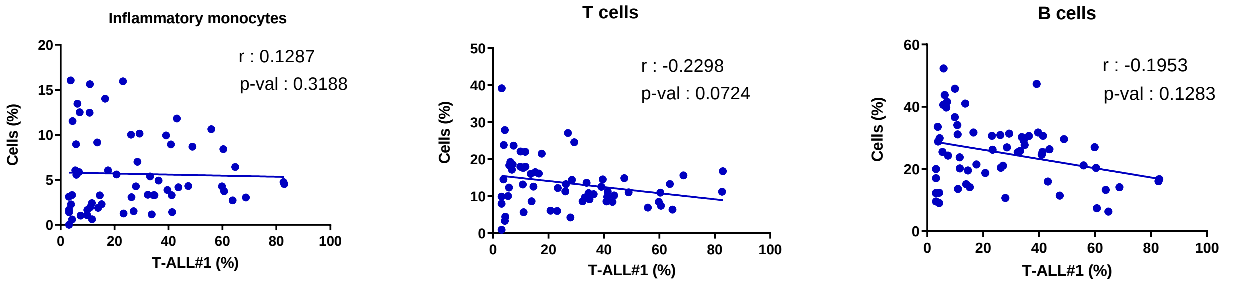


**d**

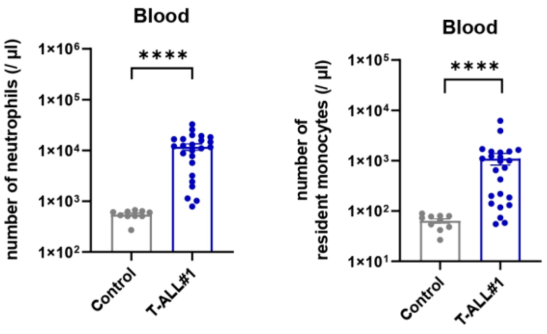

#### Supplementary Fig. S1

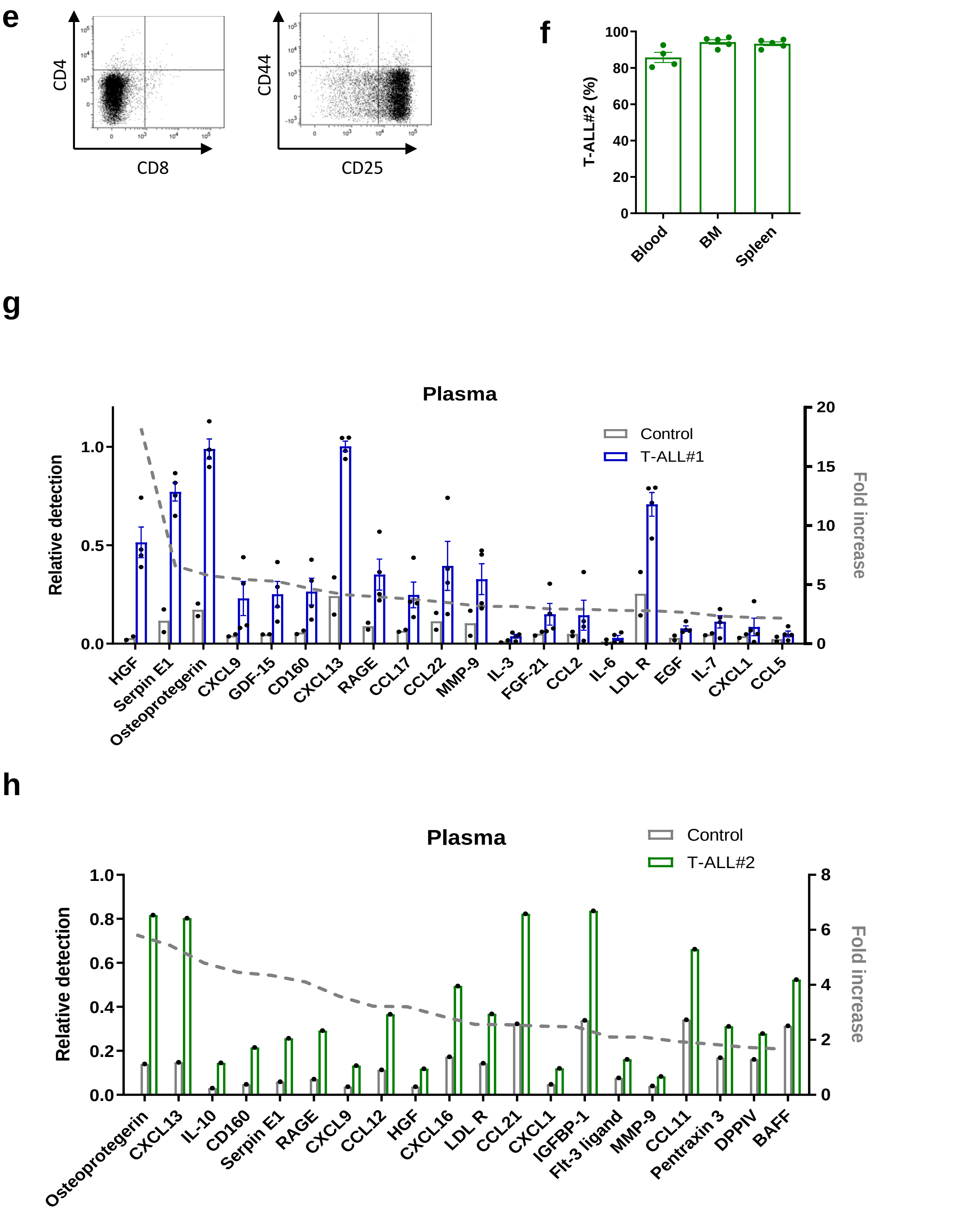

### Supplementary Fig. S2

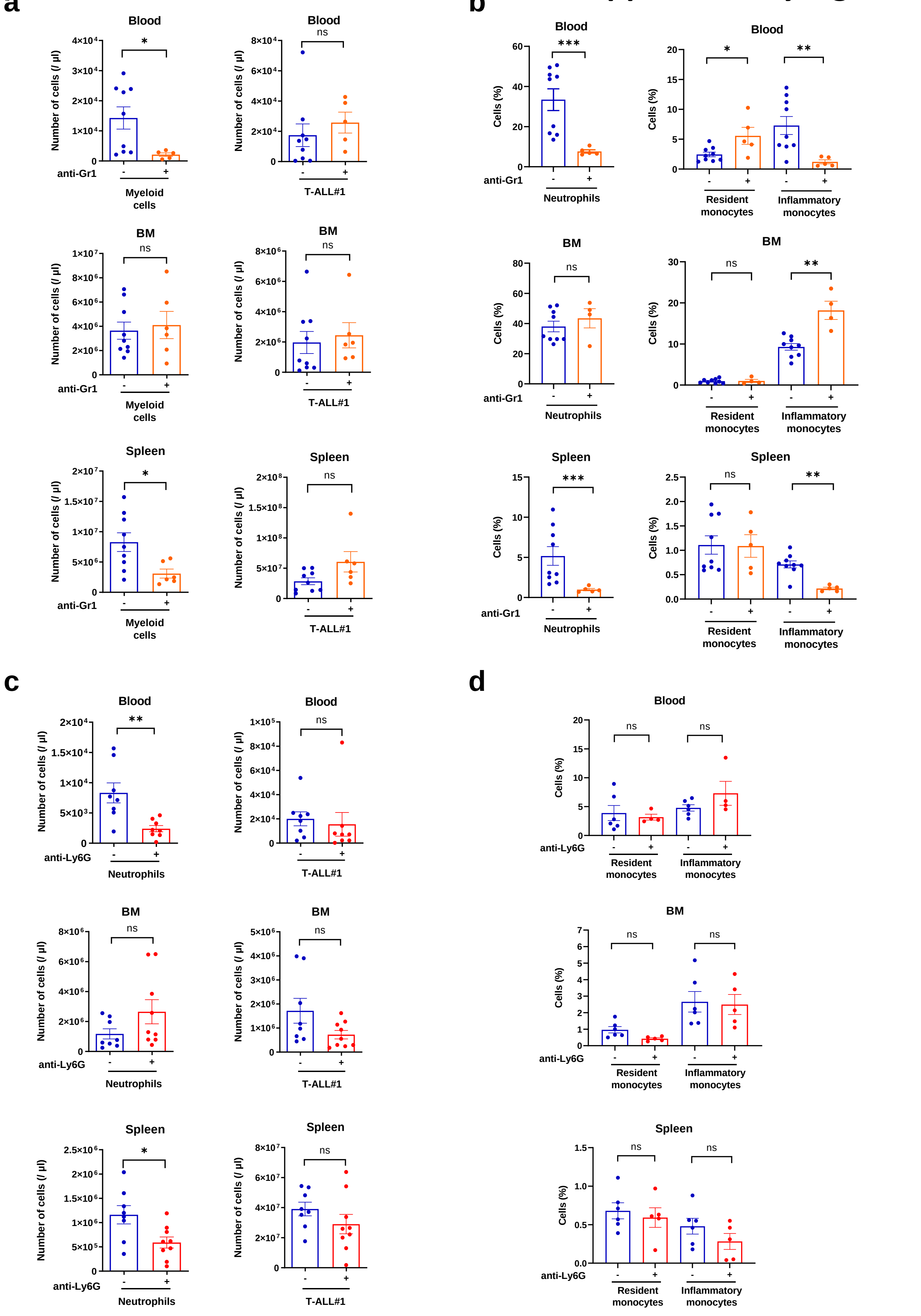

e

### Supplementary Fig. S2

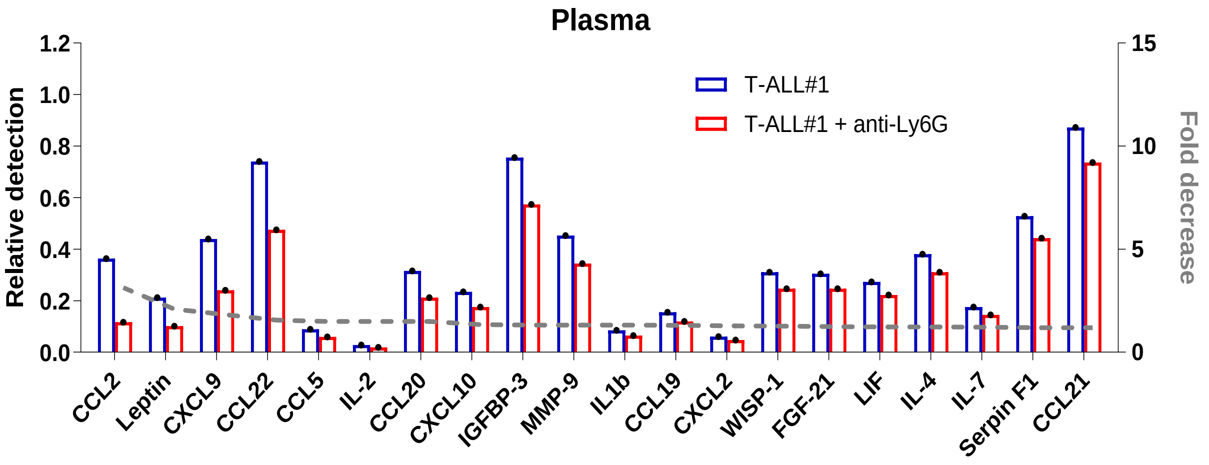

### Supplementary Fig. S3

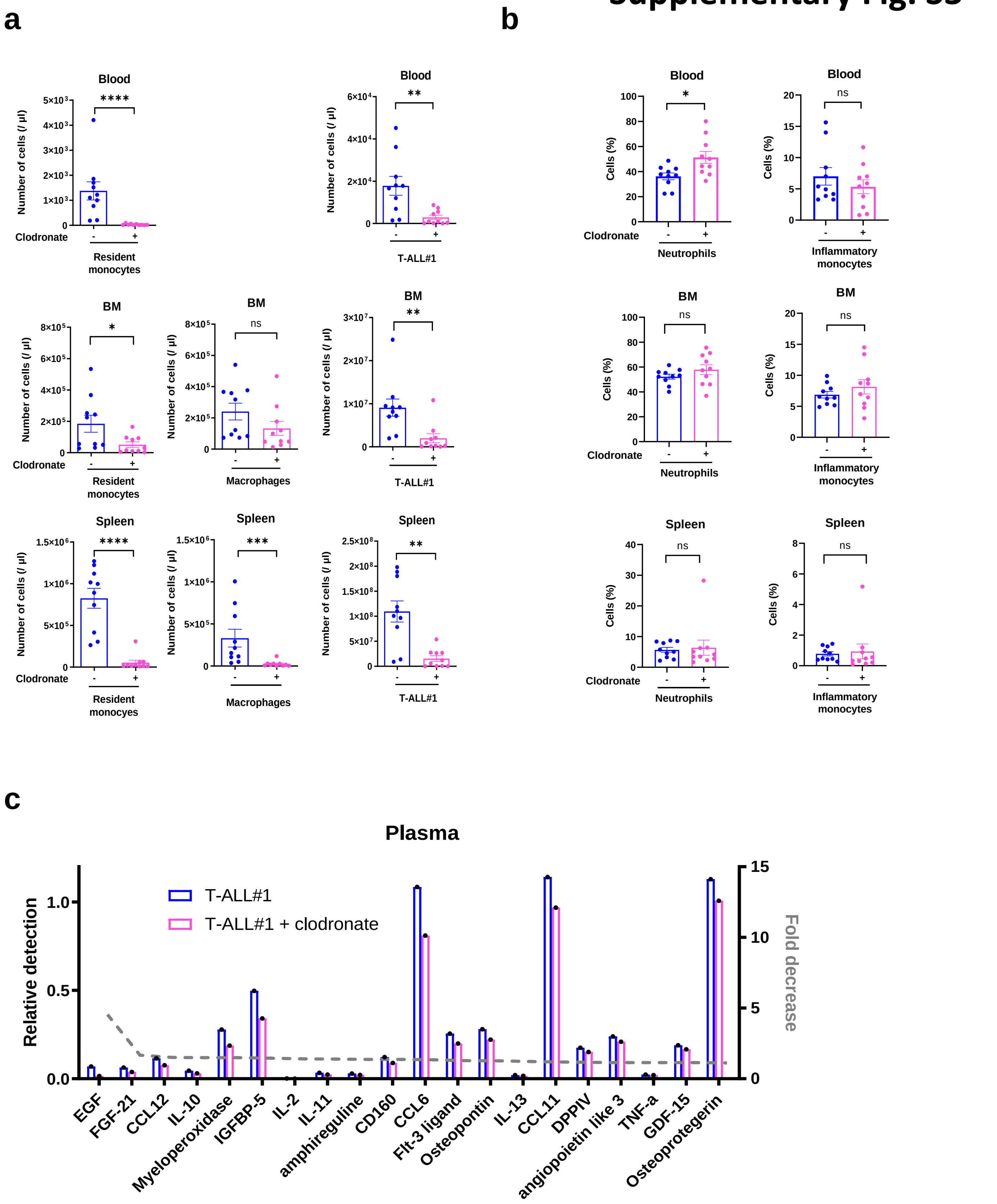

### Supplementary Fig. S4

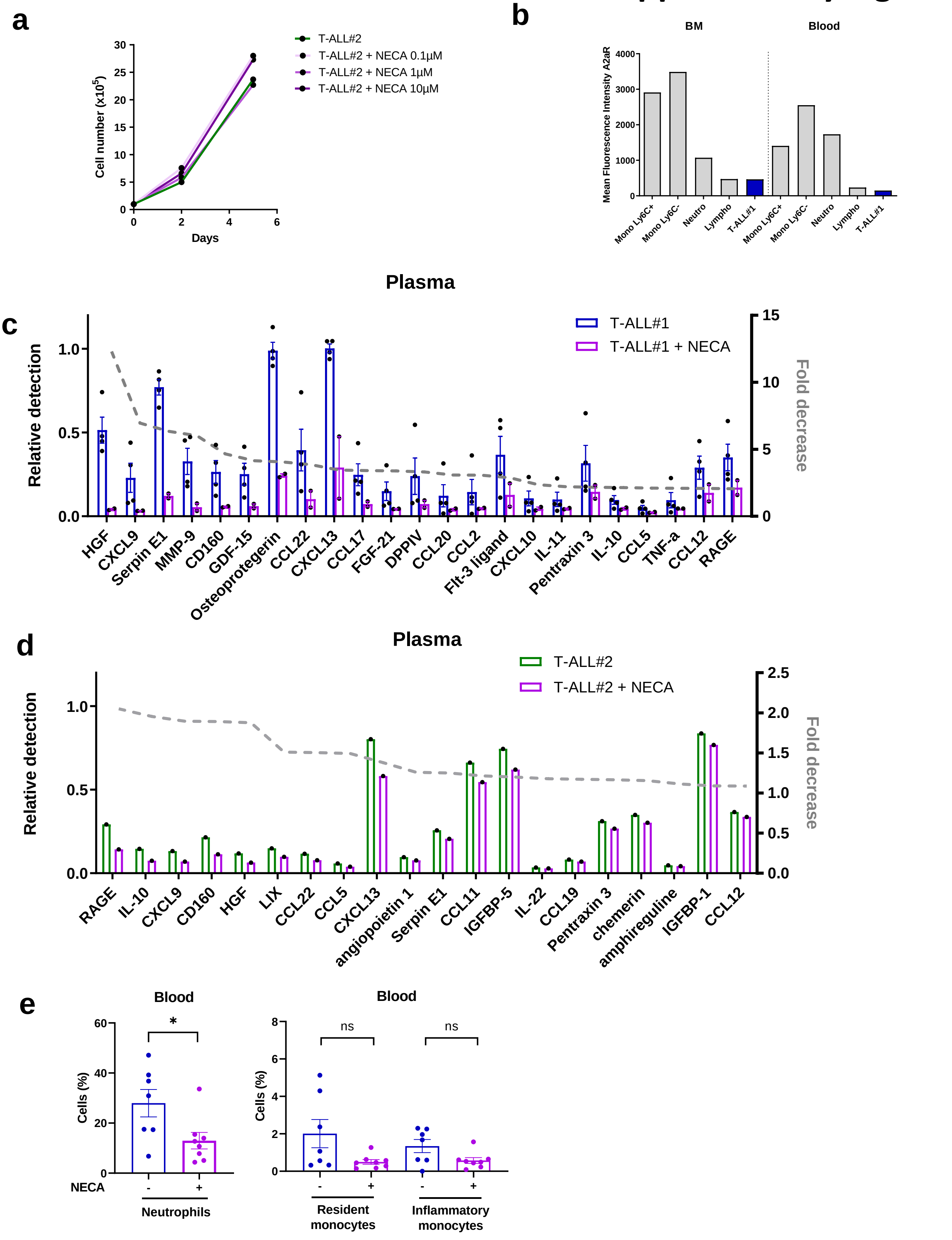

### Supplementary Fig. S5

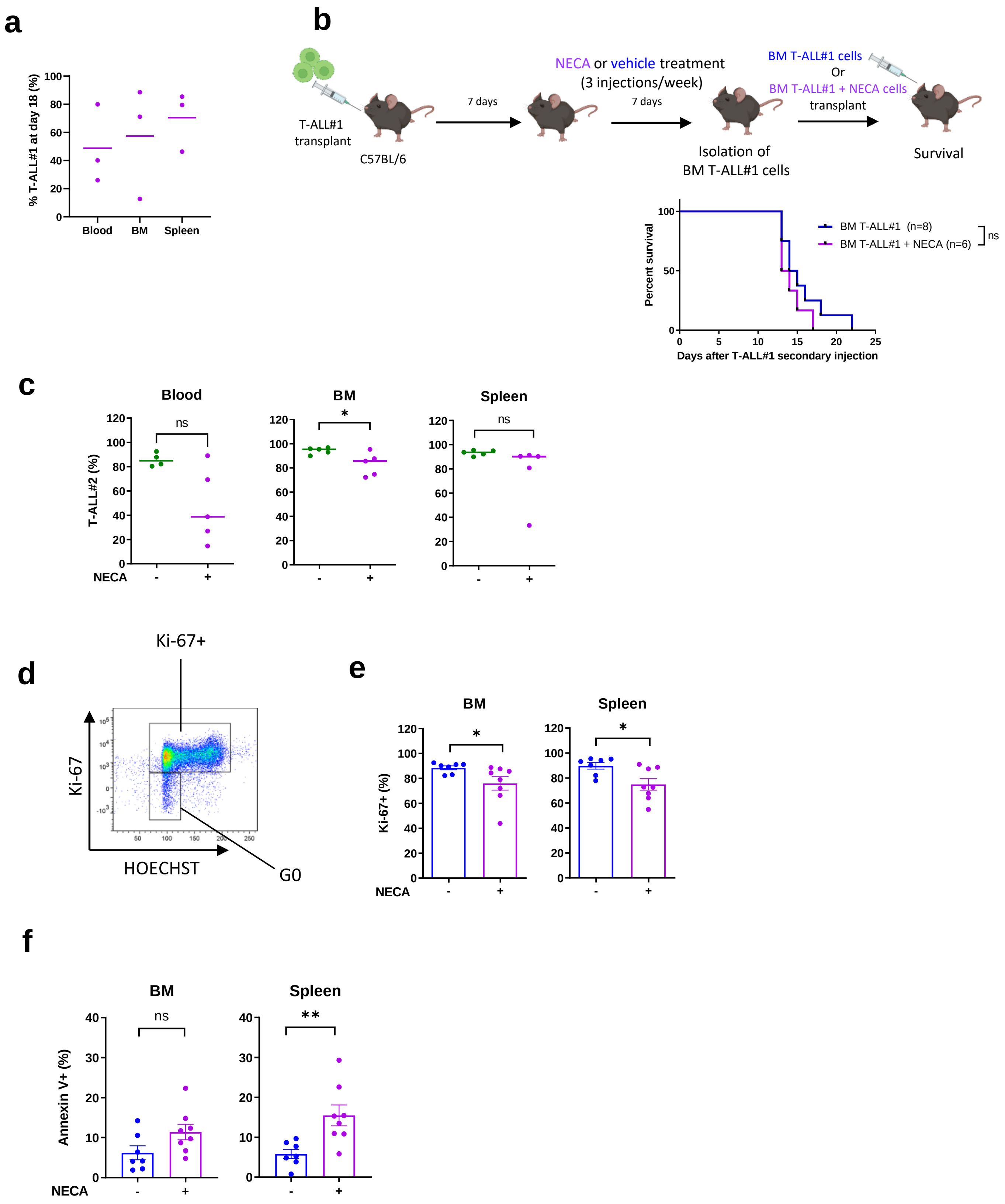

### Supplementary Fig. S6

| Ingenuity Canonical Pathways | p-value | B-H p-value | z-score |
|---|---|---|---|
| LXR/RXR Activation | 1,15E-03 | 1,91E-02 | -2,449 |
| PPAR Signaling | 5,62E-04 | 1,12E-02 | -1,633 |
| Senescence Pathway | 7,76E-03 | 4,90E-02 | -1,134 |
| Cell Cycle: G1/S Checkpoint Regulation | 4,79E-04 | 1,12E-02 | -1 |
| Melanoma Signaling | 1,29E-03 | 1,91E-02 | -1 |
| GADD45 Signaling | 2,57E-03 | 2,82E-02 | -1 |
| Th1 Pathway | 6,31E-03 | 4,47E-02 | -1 |
| Antiproliferative Role of Somatostatin Receptor 2 | 6,31E-03 | 4,47E-02 | -1 |
| JAK/STAT Signaling | 7,76E-03 | 4,90E-02 | -1 |
| PD-1, PD-L1 cancer immunotherapy pathway | 5,37E-04 | 1,12E-02 | -0,816 |
| PTEN Signaling | 5,89E-04 | 1,15E-02 | -0,447 |
| Glioblastoma Multiforme Signaling | 1,29E-03 | 1,91E-02 | -0,447 |
| HGF Signaling | 1,66E-03 | 2,14E-02 | -0,447 |
| ID1 Signaling Pathway | 3,09E-03 | 3,31E-02 | -0,378 |
| Coronavirus Pathogenesis Pathway | 3,39E-03 | 3,47E-02 | -0,378 |
| HIF1α Signaling | 3,80E-03 | 3,63E-02 | 0,378 |
| Pancreatic Adenocarcinoma Signaling | 2,04E-04 | 5,62E-03 | 0,447 |
| PI3K/AKT Signaling | 1,26E-04 | 4,07E-03 | 0,816 |
| IL-6 Signaling | 1,41E-03 | 2,04E-02 | 0,816 |
| Aryl Hydrocarbon Receptor Signaling | 4,17E-03 | 3,80E-02 | 0,816 |
| Tumor Microenvironment Pathway | 7,41E-03 | 4,90E-02 | 0,816 |
| Cyclins and Cell Cycle Regulation | 1,23E-03 | 1,91E-02 | 1 |
| IL-7 Signaling Pathway | 6,46E-03 | 4,57E-02 | 1 |
| Estrogen-mediated S-phase Entry | 4,17E-06 | 2,63E-04 | 1,342 |
| Cholecystokinin/Gastrin-mediated Signaling | 5,62E-03 | 4,27E-02 | 1,342 |
| IL-17 Signaling | 4,37E-04 | 1,12E-02 | 1,414 |
| HER-2 Signaling in Breast Cancer | 1,51E-03 | 2,09E-02 | 1,414 |
| Acute Phase Response Signaling | 2,00E-03 | 2,34E-02 | 1,633 |
| Cell Cycle Control of Chromosomal Replication | 1,91E-04 | 5,62E-03 | 2,236 |

### Supplementary Fig. S7

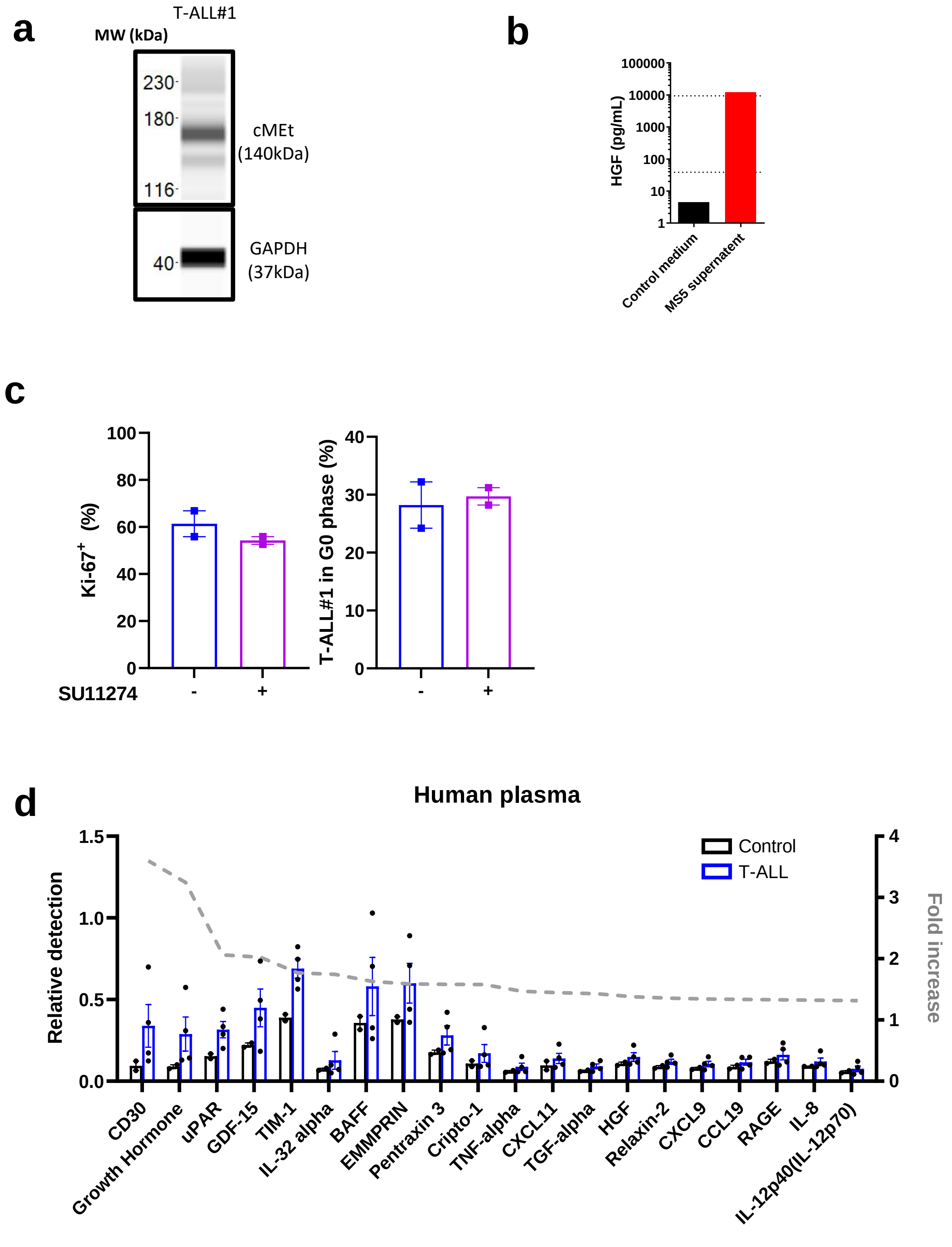

### Supplementary Table S1

| T-ALL samples | Gender | Age | Genetic abnormalities |
|---|---|---|---|
| hT-ALL#1 | M | 14 | TLX3 |
| hT-ALL#2 | M | 10 | ND |
| hT-ALL#3 | F | 6 | ND |
| hT-ALL#4 | M | 5 | t(7;10) |
| hT-ALL#5 | M | 16 | TLX3 |
| hT-ALL#6 | M | 10 | NOTCH1 mutated |
| hT-ALL#7 | M | 16 | ND |

### Supplementary Table S2

| | Primers for TAL1 and LMO1 constructions | |
|---|---|---|
| | Forward | Reverse |
| TAL1 transgene | 5’-CACAGGGTCCTTGCCAGTC-3’ | 5’-CTACCCTGCAAACAGACCTC-3’ |
| LMO1 transgene | 5’-CGGGATCCGAGATGATGGTGCTGG-3’ | 5’-CCCAAGCTTACTGAACTTGGCATTCAAAGG-3’ |

### Supplementary Table S3

| Antigen | Clone | Source |
|---|---|---|
| B220-V500 | RA3-6B2 | BD Bioscience |
| CD115-PE | ASF98 | BioLegend |
| CD115-PE/Cyanine 7 | ASF98 | BioLegend |
| CD11b-BV605 | M1/70 | BioLegend |
| CD11b-PECyanine 7 | M1/70 | BioLegend |
| CD25-BV421 | PC61 | BioLegend |
| CD3-PerCP/Cyanine5.5 | 145-2C11 | BioLegend |
| CD44-APC/Cyanine 7 | IM7 | BioLegend |
| CD45.1-APC | A20 | BioLegend |
| CD45.2-FITC | 104 | BioLegend |
| CD4-BV421 | GK1.5 | BioLegend |
| CD4-PerCP/Cyanine 5.5 | GK1.5 | BioLegend |
| CD8a-APC | 53-6.7 | eBioscience |
| CD8a-PE | 53-6.7 | BD pharmingen |
| cMet-PE | eBioclone 7 | Invitrogen eBioscience |
| Phospho cMet-APC | MetY12341235-6F11 | Invitrogen |
| F4/80-AF700 | BM8 | BioLegend |
| Gr1-PerCP/Cyanine 5.5 | RB6-8C5 | BioLegend |
| Ly6C-APC | HK1.4 | Invitrogen |
| Ly6C-BV711 | HK1.4 | BioLegend |
| Ly6G-BV421 | 1A8 | BioLegend |

### Supplementary Table S4

| Antigen | Clone | Reference | Supplier |
|---|---|---|---|
| cMet | E4E4W | 82202S | Cell Signaling Technology |
| GAPDH | 14C10 | 2118 | Cell Signaling Technology |
